# CLOVER: Chemical-gene LLM benchmark for obtaining valid entity relationships for the Comparative Toxicogenomics Database (CTD)

**DOI:** 10.64898/2026.11.10.765703

**Authors:** Sanjana Cheerla, Allan Peter Davis, Thomas C. Wiegers, Steffen Heber, Carolyn J. Mattingly, Munindar P. Singh

## Abstract

**Objective:** The Comparative Toxicogenomics Database (CTD) provides expert-curated chemical-gene interactions about environmental exposure effects on human diseases. Existing automated relation extraction systems are typically restricted to 3-10 interaction types and operate on abstracts only. We introduce Clover, a systematic benchmark for evaluating large language models (LLMs) on full-text chemical-gene interaction extraction using CTD-defined chemical–gene interaction terminology. The framework is designed to support 47 base interactions with directional modifiers (increase, decrease, affects), yielding 136 possible supported interaction classes. Evaluation is restricted to the subset of classes represented in the selected articles.

**Methods:** Clover processes 50 full-text PubMed Central articles (1,563 sections) with PubTator and CTD integration. The pipeline consists of four stages: (1) LLM Extraction with one-shot and few-shot prompting; (2) LLM Validation to confirm and revise interactions; (3) Rule-based Refinement of extracted interactions to standardize terminology and eliminate errors; and (4) Human Review for expert oversight. We benchmark four LLMs (GPT-4.1, GPT-4o, Claude-Sonnet-4.5, and DeepSeek-V4.1-Flash) and compare against a naive baseline that applies unstructured prompting directly to full-text PDFs.

**Results:** The naive baseline achieves only 13% recall and 36% precision. Refinement significantly enhances performance by eliminating unsupported interactions. Under strict limitations, Claude-Sonnet-4.5 alone achieves 72% recall and 61% precision, and Claude with self-validation achieves 74% recall and 49% precision compared to CTD. Of the 136 supported interaction classes, 17 were represented in our evaluation.

**Conclusion:** To our knowledge, Clover establishes the first benchmark for full-text chemical-gene interaction extraction aligned with CTD. Our findings demonstrate that Clover achieves workable metrics for CTD-aligned curation relative to minimal whole-PDF prompting. This study addresses only a fraction of the content manually curated in CTD; considerably more work is needed to advance LLM-based tools to determine the extent to which these tools can approximate the full capacity of manual curation at CTD.

## Introduction

The Comparative Toxicogenomics Database (CTD; https://ctdbase.org/) [1] is a valuable resource in biomedical research, providing manually curated data describing complex chemical-gene-phenotype-disease interactions and exposure conditions. CTD primarily relies on manual curation from the scientific literature, where domain experts review and extract relevant interactions from biomedical texts using community-based and standardized vocabularies and ontologies. Manual curation is inherently labor-intensive and constrained by the availability of domain experts [2].

Furthermore, variability can exist in how authors convey entity interactions in biomedical articles, potentially complicating manual curation consistency. Although CTD has conducted extensive quality assurance studies [3] on its curation, a reliable, reproducible, and precise automated approach could help further address these challenges for manual curation processes by enhancing consistency and efficiency in extracting meaningful interactions. To test this hypothesis, we use Large Language Models (LLMs) to extract interactions between chemicals and genes described in biomedical texts according to CTD curation policies. Specifically, we implement a systematic benchmark, Clover, to support 47 interactions (S2 Fig 1) where most interaction types can be prefixed with increase, decrease, or affect, yielding 136 unique chemical-gene interaction types. In practice, full-text articles exhibit an imbalanced distribution concentrated in a subset of these interactions, and our evaluation covers a subset of these interactions.

**Fig 1.**
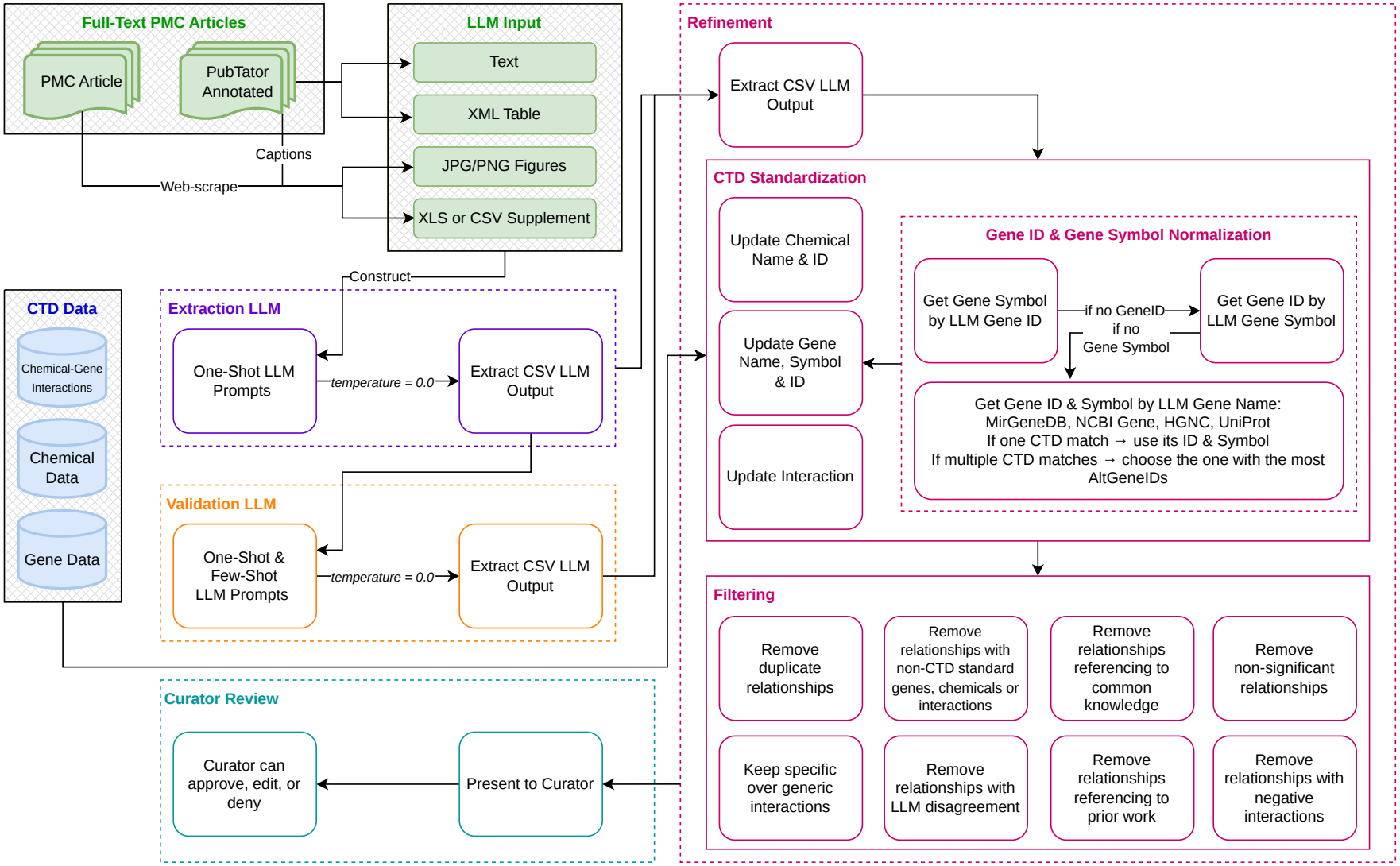
Overview of the Clover pipeline. First, PubTator-annotated PubMed articles (green) and CTD sources (indigo) are processed to extract chemical-gene interactions using chained Large Language Models (LLMs). The pipeline involves four stages: 1) (purple) Extraction LLM, where one-shot and few-shot LLM responses are generated to extract potential interactions; 2) (orange) Validation LLM, where the output of the Extraction LLM is validated using few-shot prompting; 3) (magenta) Refinement Validation to comply with CTD standards; 4) (teal) Curator Review, where an expert reviews the interactions and approves, edits, or denies them. This pipeline is not implemented in CTD.

Biomedical relation extraction (BioRE) is essential because it converts raw data into organized biomedical knowledge. This structured data supports future research in drug discovery, clinical therapies, pathological studies, and more, allowing researchers to make data-driven advances [4]. Challenges in BioRE include handling the vast amount of available literature, the variability in terminology and language use, and the need for high performance in extracting meaningful interactions.

Relation extraction (RE) is a specific Natural Language Processing (NLP) task. Early studies of RE methods include rule-based and traditional machine learning approaches [5]. Rule-based models use annotated rules and patterns to find interactions [6]. Traditional machine learning techniques involve classification tasks to extract sentences containing interactions [7]. However, these methods depend on the dataset and require extensive domain-specific expertise to extract interactions between biomedical entities. Also, the variability in biomedical literature often limits these approaches.

Transformer-based models such as BERT [8] and its pre-trained biomedical variant, BioBERT [9], have shown promising results. However, these often require extensive fine-tuning with domain-specific datasets, which can be time-consuming due to the need for large annotated data [10].

More recently, LLMs have gained attention in biomedical research for their ability to process complex data. Asada and Fukuda [11] propose a BioRE method, showing that few-shot learning performs poorly in biomedical tasks and LLM-generated entity explanations improve classification-based extraction. Zhou et al. [12] introduce LEAP, a framework for adapting LLMs to BioRE tasks through instruction-based and example-driven prompting for zero-shot, one-shot, and few-shot scenarios. One-shot learning involves providing exactly one example to guide the LLM, distinguishing it from zero-shot learning, which uses no prior examples, and few-shot learning, which uses a small number of examples.

Prior studies have evaluated LLMs on biomedical relation extraction benchmarks; earlier works typically restrict relation classification to fewer than ten classes and focus on abstracts or sentence-level annotations [11, 12]. For instance, Zhang et al. introduce ChemDisGene [13], a CTD-derived benchmark for extracting relationships among chemicals, diseases, and genes from abstracts. More recently, LLM-based approaches have extended to document-level extraction and end-to-end evaluation [14, 15]. However, systematic evaluation of CTD-aligned chemical–gene interactions across full-text article components, including tables, figures, and supplements, remains limited.

Since its initial release in 2004, CTD has relied on manual curation, supported in part by existing text-mining tools such as PubTator [16], which provides automated named entity recognition (NER) for key biomedical concepts. PubTator facilitates curation by identifying and normalizing mentions of chemicals, genes, and diseases in the literature. Building on this foundation, Clover extends the pipeline by applying LLMs to extract chemical-gene interactions from full-text articles.

A central component of this work is understanding where LLMs succeed and fail. We use LLMs to curate chemical-gene interactions across different PubMed Central [17] full-text article sections by key content-rich sections, including results, figures, tables, and CSV-formatted supplements. We further investigate whether chaining LLMs, where a second model validates the output of the first, improves extraction quality.

The key contributions of this work are:

1. **Benchmarking LLMs for BioRE:** We provide a practical, systematic benchmark of four state-of-the-art general-purpose LLMs—GPT-4.1, GPT-4o, DeepSeek-V4.1-Flash, and Claude-Sonnet-4.5—across single-model and chained configurations for chemical-gene CTD-aligned relation extraction using PubTator for named entity recognition of chemicals and genes. Performance is evaluated across various full-text PMC article section types (main text, tables, figures, and CSV- and XLS-styled supplements).
2. **Baseline Comparison:** We compare structured, detailed one-shot and few-shot prompts against a minimal zero-shot baseline that processes full-text PDFs without section-level preprocessing or PubTator annotations, providing a practical reference for evaluating performance of Clover.
3. **Evaluating LLM Chaining:** We systematically compare single-model and chained LLM configurations. We find that chaining does not consistently improve performance, and only three chained configurations improve precision and/or recall. Standalone Claude-Sonnet-4.5 achieves the highest precision and one of the highest recall values for BioRE. To the best of our knowledge, no prior study has evaluated chained LLM configurations for chemical-gene BioRE with CTD-aligned interaction vocabulary.
4. **Developing the C****lover** **Approach:** Clover provides an end-to-end human-review pipeline for extracting and refining chemical-gene interactions from biomedical literature to comply with CTD standards. Extracted interactions can then be reviewed and validated by domain experts.

This study sheds light on the potential and limitations of general-purpose LLMs for BioRE tasks. Our findings highlight differences in LLM performance, the limitations of LLM chaining, and actionable guidance for developing future LLM-based biomedical curation systems.

## Background on LLMs

Large Language Models (LLMs) are trained on massive textual corpora, enabling them to perform advanced language tasks with minimal task-specific supervision. In biomedical NLP, they are increasingly applied across both tasks and domains, with a reduced reliance on large annotated datasets.

### Single LLMs for Zero, One, and Few-Shot BioRE

LLMs can perform relation extraction using one-, few-, and zero-shot learning, achieving state-of-the-art (SOTA) performance without task-specific training [18]. For example, GPT-3.5 and GPT-4 have been tested on sentence-based chemical-disease or gene-disease relation extraction in zero- and one-shot settings, resulting in F-1 scores of around 0.8 on benchmarks like EU-ADR (a corpus of drug-adverse event associations), GAD (Genetic Association Database, containing gene-disease associations), and ChemProt (a dataset of chemical-protein interactions) [19]. In contrast, we apply LLMs to full-text articles. We extract multiple chemical-gene interactions from complex, unstructured contexts. This setting is significantly more complex than sentence-level tasks.

Asada and Fukuda [11] incorporate an in-context few-shot RE model using LLM prompts and find that naïve few-shot prompting alone underperforms on specialized biomedical RE datasets, and that including additional domain context improves performance. Similarly, Zhao et al. [20] propose a prompt-based RE method using a T5 transformer with a masked prompt template plus K-nearest neighbors retrieval of examples for few-shot learning for document-level BioRE.

Prior works have used a single LLM to identify interactions from text via prompt-based generation. Choudhury and Dey [21] present a GPT-3-based information extraction approach where the LLM is given a few in-context examples and extracts relevant entities and interactions from unstructured text, achieving, at best, moderate precision and recall. Chen et al. [22] find that closed-source LLMs (like GPT-3/GPT-4) are outperformed by SOTA fine-tuning approaches in most cases, including relation extraction. Gill et al. [23] use LLMs to achieve high performance on gene–protein interactions from abstracts, which they noted was less complex than automatically extracting pieces of text from the body of the paper.

Recent studies have extended LLM-based BioRE beyond sentence-level tasks. Dong et al. [14] introduce SyRACT, which combines retrieval-augmented generation and chain-of-thought reasoning for document-level biomedical relation extraction. Brokman et al. [15] evaluate end-to-end zero-shot extraction using LLMs on seven biomedical datasets.

Laskar et al. [24] investigate LLM-based evaluation of BioRE, highlighting the importance of structured output formats. More broadly, LLMAgent4Bio [25] surveys integration of LLMs with external tools, biological knowledge, and multistep reasoning. Building on these directions, Clover focuses specifically on CTD-aligned chemical–gene interaction extraction from full-text PMC article components, using a controlled interaction vocabulary and evaluating agreement with CTD-curated relationships.

These studies demonstrate that LLMs can perform BioRE with minimal training data. However, they typically evaluate performance on datasets containing only a few types of interactions, and on sentences and/or abstracts rather than full text. Unlike prior studies, Clover supports a broader CTD interaction vocabulary, although only a subset of these classes is represented in our evaluation. Additionally, LLMs lack task-specific training, and in complex cases, more advanced prompting and additional feedback may be needed.

### Chained LLMs for Information Extraction

Table 1 summarizes key differences between Clover and prior BioRE works, highlighting that Clover is the only approach to evaluate LLMs on full-text articles and supplementary materials aligned with CTD controlled vocabulary in a chained LLM setting.

**Table 1.**
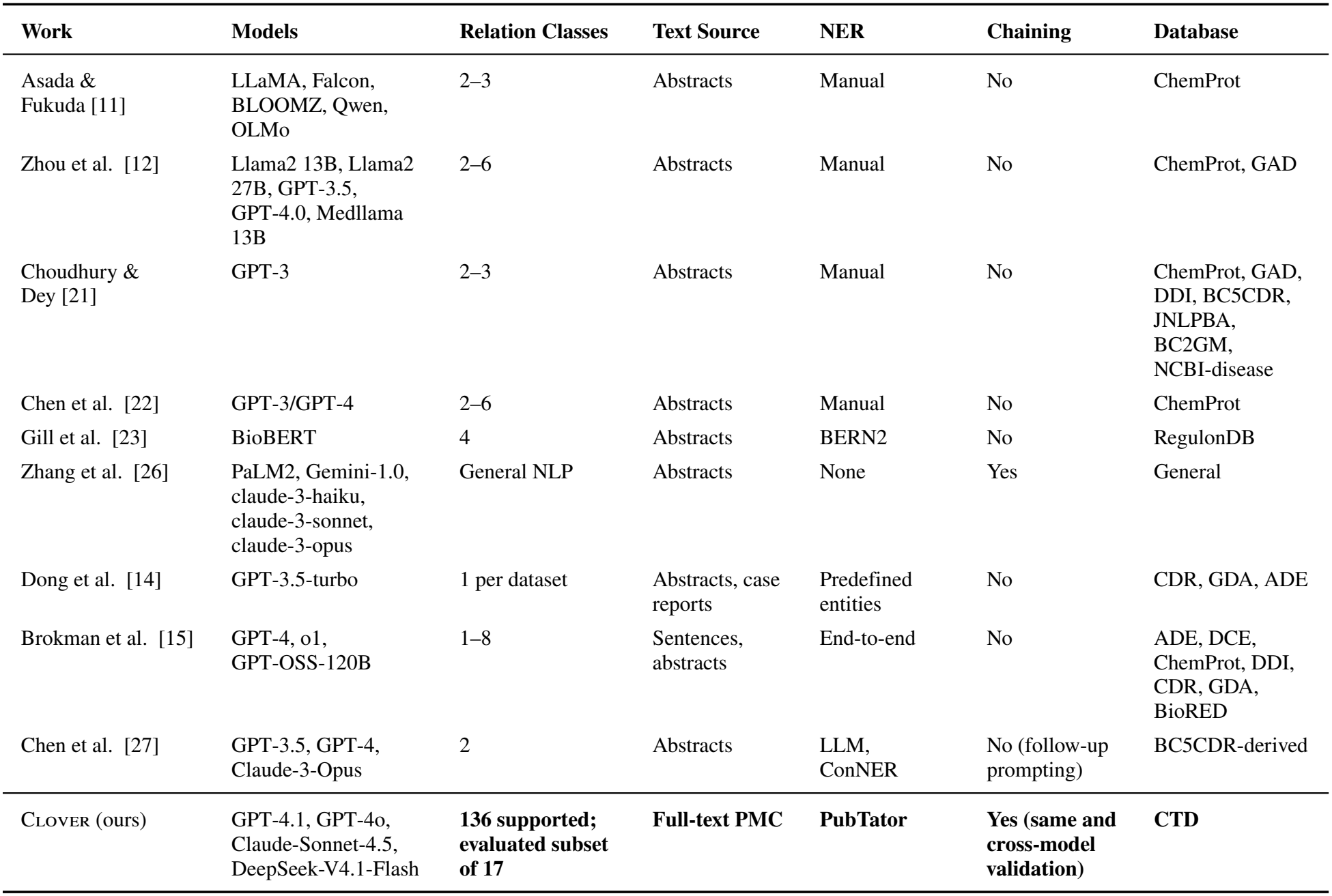
Comparison of Clover with previous biomedical relation extraction works.

| Work | Models | Relation Classes | Text Source | NER | Chaining | Database |
|---|---|---|---|---|---|---|
| Asada & Fukuda [11] | LLaMA, Falcon, BLOOMZ, Qwen, OLMo | 2–3 | Abstracts | Manual | No | ChemProt |
| Zhou et al. [12] | Llama2 13B, Llama2 27B, GPT-3.5, GPT-4.0, Medllama 13B | 2–6 | Abstracts | Manual | No | ChemProt, GAD |
| Choudhury & Dey [21] | GPT-3 | 2–3 | Abstracts | Manual | No | ChemProt, GAD, DDI, BC5CDR, JNLPBA, BC2GM, NCBI-disease |
| Chen et al. [22] | GPT-3/GPT-4 | 2–6 | Abstracts | Manual | No | ChemProt |
| Gill et al. [23] | BioBERT | 4 | Abstracts | BERN2 | No | RegulonDB |
| Zhang et al. [26] | PaLM2, Gemini-1.0, claude-3-haiku, claude-3-sonnet, claude-3-opus | General NLP | Abstracts | None | Yes | General |
| Dong et al. [14] | GPT-3.5-turbo | 1 per dataset | Abstracts, case reports | Predefined entities | No | CDR, GDA, ADE |
| Brokman et al. [15] | GPT-4, o1, GPT-OSS-120B | 1–8 | Sentences, abstracts | End-to-end | No | ADE, DCE, ChemProt, DDI, CDR, GDA, BioRED |
| Chen et al. [27] | GPT-3.5, GPT-4, Claude-3-Opus | 2 | Abstracts | LLM, ConNER | No (follow-up prompting) | BC5CDR-derived |
| Clover (ours) | GPT-4.1, GPT-4o, Claude-Sonnet-4.5, DeepSeek-V4.1-Flash | **136 supported; evaluated subset of 17** | **Full-text PMC** | **PubTator** | **Yes (same and cross-model validation)** | **CTD** |

Emerging research explores chained LLM architectures where multiple LLMs interact sequentially (one validating or refining the output of another) to improve information extraction. Grunde-McLaughlin et al. [28] survey over 100 such approaches to form a design space for LLM chains, stating that chaining can decompose complex tasks into subtasks and use later LLMs to catch errors made by earlier ones. Zhang et al. [26] introduce a framework where several LLMs collaborate on a task, with each agent working on a portion of the task and passing its results to another LLM that synthesizes the final answer. This chained LLM approach is used to handle long texts, and it significantly outperforms single-model baselines [26].

The LLM chain framework can be applied to relation extraction, where an initial LLM identifies candidate interactions in text, and a second LLM reviews them for correctness and consistency before finalizing the output. Although research specific to chained LLMs for BioRE is still nascent, general NLP studies indicate that using chained LLMs can increase reliability, reduce errors, and curb hallucinations to ensure factual accuracy in extracted information [28]. Chen et al. [27] explore a follow-up LLM approach for document-level chemical-disease relation extraction, where the same model is prompted to further refine extracted relationships.

Clover differs from previous works by supporting extraction across a greater number of more complex interactions. Additionally, we systematically evaluate four models individually and cross-model extractor–validator configurations to assess whether a second LLM improves extraction performance. Unlike prior methods, our pipeline processes biomedical literature with minimal domain-specific training by leveraging general-purpose LLMs. Furthermore, we construct highly structured CSV-style responses that include chemical and gene names and IDs, interaction type, and source text, enabling downstream integration into CTD.

## Materials and methods

Our approach (Fig. 1) leverages PubTator [16] for named entity recognition of chemicals and genes. We then use LLMs to automate the extraction of chemical-gene interactions from biomedical texts, potentially reducing the time and effort required for manual curation.

### Data Sources

We randomly selected 50 full-text papers using PubMed IDs that are the source for CTD chemical-gene interaction data. We mapped each CTD PubMed ID to its corresponding PubMed Central ID to identify articles with available full text. We used PubTator to download chemical- and gene-annotated versions of the 50 full-text articles. To download figures, XLS, and/or CSV supplement data, we built a web scraper.

We then separated each article into individual data components as shown in Fig. 1:

1. **Text:** Chemical and PubTator-annotated text paragraphs.
2. **Table:** XML table and captions.
3. **Figures:** JPG/PNG figures and captions.
4. **Supplement:** CSV or XLS supplemental data, captions, and XLS sheet names if available.

Articles were randomly selected from CTD-associated PMIDs without stratification by research topic, chemical class, organism, experimental setting, or number of CTD-curated interactions. The only eligibility check was whether the selected PMID corresponded to a full-text article available in PMC with PubTator full-text annotations. Therefore, the final sample is restricted to articles and is not necessarily representative of the broader CTD-curated literature. We selected a total of 50 full-text PMC articles containing 1,563 processed data components: 981 text sections, 77 tables, 309 figures, and 196 supplementary components (Table 2). Four articles contained CSV-based supplementary tables. A list of articles, CTD-curated relationships, and data component details can be found in Table 1 in S6.

**Table 2.**
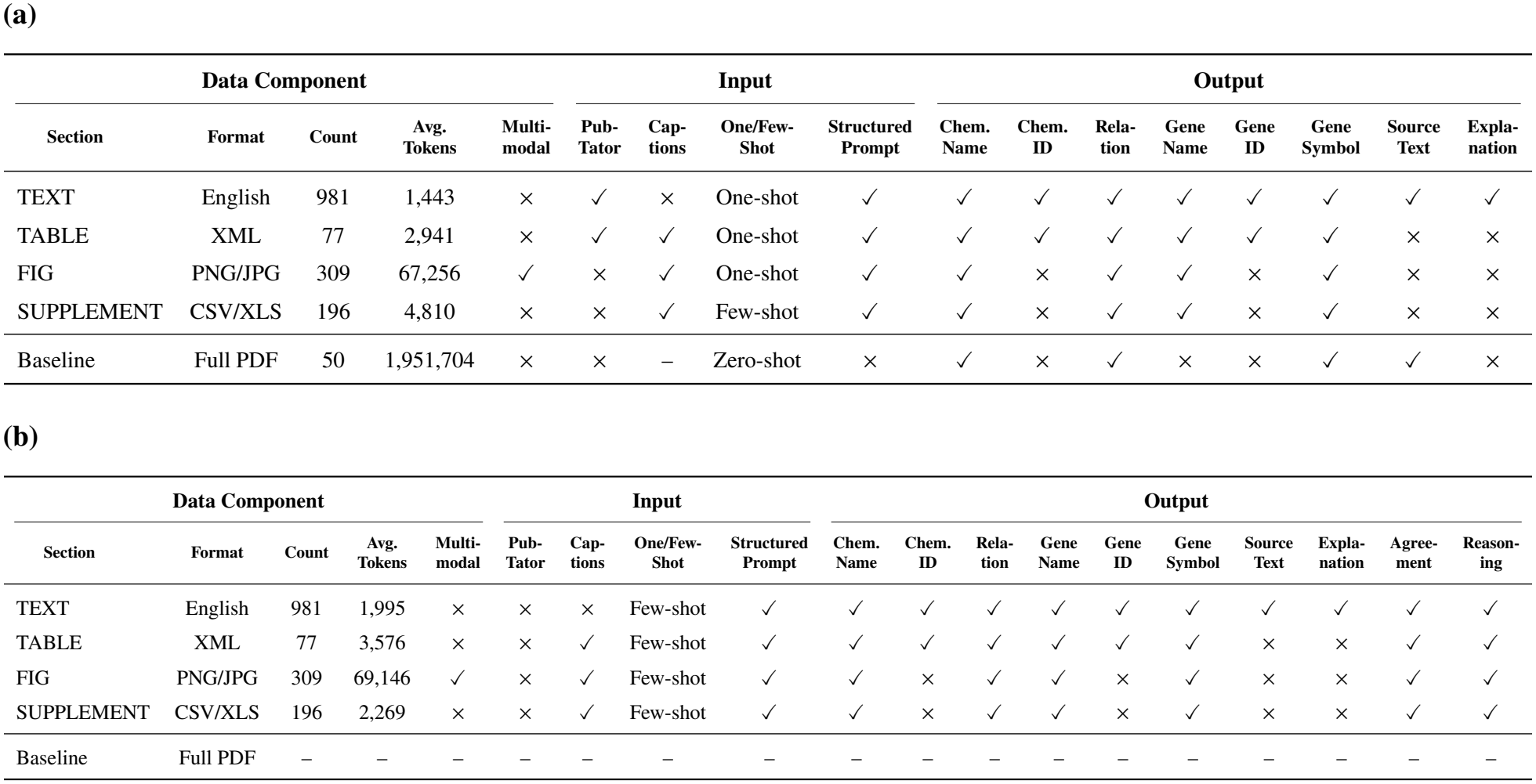
Summary of data components, LLM inputs, and outputs used in Clover extraction (Table 2a) and validation (Table 2b) prompts. *Output* shows fields present for each section type; *Count* is number of prompts, *Avg. Tokens* is the prompt length. *Multimodal* indicates text only or inclusion of encoded images. PubTator provides only chemical and gene entity classification. *Source Text* and *Explanation* in TEXT sections support traceability with verbatim evidence.

| **Data Component** |  |  |  |  | **Input** |  |  |  | **Output** |  |  |  |  |  |  |  |  |  |
|---|---|---|---|---|---|---|---|---|---|---|---|---|---|---|---|---|---|---|
| **Section** | **Format** | **Count** | **Avg. Tokens** | **Multi-modal** | **Pub-Tator** | **Cap-tions** | **One/Few-Shot** | **Structured Prompt** | **Chem. Name** | **Chem. ID** | **Rela-tion** | **Gene Name** | **Gene ID** | **Gene Symbol** | **Source Text** | **Expla-nation** |  |  |
| TEXT | English | 981 | 1,443 | × | ✓ | × | One-shot | ✓ | ✓ | ✓ | ✓ | ✓ | ✓ | ✓ | ✓ | ✓ |  |  |
| TABLE | XML | 77 | 2,941 | × | ✓ | ✓ | One-shot | ✓ | ✓ | ✓ | ✓ | ✓ | ✓ | ✓ | × | × |  |  |
| FIG | PNG/JPG | 309 | 67,256 | ✓ | × | ✓ | One-shot | ✓ | ✓ | × | ✓ | ✓ | × | ✓ | × | × |  |  |
| SUPPLEMENT | CSV/XLS | 196 | 4,810 | × | × | ✓ | Few-shot | ✓ | ✓ | × | ✓ | ✓ | × | ✓ | × | × |  |  |
| Baseline | Full PDF | 50 | 1,951,704 | × | × | – | Zero-shot | × | ✓ | × | ✓ | × | × | ✓ | ✓ | × |  |  |
| **(b)** |  |  |  |  |  |  |  |  |  |  |  |  |  |  |  |  |  |  |
| **Data Component** |  |  |  |  | **Input** |  |  |  | **Output** |  |  |  |  |  |  |  |  |  |
| **Section** | **Format** | **Count** | **Avg. Tokens** | **Multi-modal** | **Pub-Tator** | **Cap-tions** | **One/Few-Shot** | **Structured Prompt** | **Chem. Name** | **Chem. ID** | **Rela-tion** | **Gene Name** | **Gene ID** | **Gene Symbol** | **Source Text** | **Expla-nation** | **Agree-ment** | **Reason-ing** |
| TEXT | English | 981 | 1,995 | × | × | × | Few-shot | ✓ | ✓ | ✓ | ✓ | ✓ | ✓ | ✓ | ✓ | ✓ | ✓ | ✓ |
| TABLE | XML | 77 | 3,576 | × | × | ✓ | Few-shot | ✓ | ✓ | ✓ | ✓ | ✓ | ✓ | ✓ | × | × | ✓ | ✓ |
| FIG | PNG/JPG | 309 | 69,146 | ✓ | × | ✓ | Few-shot | ✓ | ✓ | × | ✓ | ✓ | × | ✓ | × | × | ✓ | ✓ |
| SUPPLEMENT | CSV/XLS | 196 | 2,269 | × | × | ✓ | Few-shot | ✓ | ✓ | × | ✓ | ✓ | × | ✓ | × | × | ✓ | ✓ |
| Baseline | Full PDF | – | – | – | – | – | – | – | – | – | – | – | – | – | – | – | – | – |

Only text and table data annotated by PubTator, when available, were provided to the LLM (Table 2). Since supplements can appear in multiple forms, we restrict them to CSV- or XLS-formatted supplements for consistency and noise reduction. Additionally, several CTD data sources are incorporated throughout Clover. Below is a description of these CTD data sources and their respective roles:

#### CTD Chemical-gene Interaction Data

We use this data to compare our results against CTD expert-curated data. CTD interaction terminology was also provided in the extraction and validation prompts to guide relationship classification.

#### CTD Relation Terminology

This data is provided to the LLM in the Extraction and Validation step to assist in selecting CTD Relationship Data. A standard CTD interaction consists of a prefix and the interaction name. There are three possible prefixes which we consider—’increase(s)’, ‘decrease(s)’, and ‘affect(s)’—and 47 different interaction names, resulting in 136 possible interaction classes (S2 Fig. 1). Of the 136 supported interaction classes, 17 were represented in the CTD reference interactions for the selected articles. An example of an interaction would be ‘increases expression’. This data is also used in Refinement to standardize terminology from LLM output.

#### CTD Chemical Data and CTD Gene Data

This data is used in Refinement to replace the LLM-extracted chemical names, chemical IDs, gene names, gene symbols, and gene IDs with their CTD standard counterparts. It includes the MeSH ID and CTD standard name for chemicals, gene symbols, and NCBI IDs for genes.

CTD stores single- and multi-step interactions. Single-step interactions involve one chemical, one gene, and one interaction. Multi-step interactions include multiple entities and multiple interactions where one reaction affects another. This study is designed to extract only single-step interactions.

Since no prior work extracts chemical-gene interactions from full-text articles using a CTD-aligned classification scheme, we compare Clover against a naive zero-shot baseline (S2 Baseline). The baseline applies a minimal unstructured prompt directly to full-text PDFs of the 50 articles.

### The Clover Approach

Fig. 1 displays a step-by-step overview of Clover. There are four major steps: Extraction LLM, Validation LLM, Refinement, and Curator Review. The pipeline from extraction to refinement is completely automated; the curator review step has been conceived as a human-review component. The pipeline is currently experimental and not yet formally implemented in CTD.

PubMed articles are processed to extract text data, XML tables, and captions containing chemicals and genes using PubTator [16] annotations. These data components, along with CTD interaction data and PubTator annotations, are used to generate one-shot and few-shot learning prompts. The prompts are then fed into the Extraction LLM step. The goal of the first LLM is to identify potential chemical-gene interactions from the passage using predefined interaction categories and information regarding chemicals and genes present in the passage.

Next, the passage and output from the Extraction LLM are sent to the Validation LLM. The goal of the second LLM is to check consistency, correctness, and alignment with CTD-standard terminology. Specifically, the LLM verifies if the extracted interactions are well-supported by the text and not from prior work using the PMC article context. The second LLM agrees or disagrees with the extracted interactions and replaces them with CTD interaction terminology if needed.

To control the behavior of LLM outputs, we selected fixed generation hyperparameters. For Extraction and Validation LLMs, we set the *temperature* hyperparameter to 0.0 for deterministic and slightly flexible responses. This ensures that validation is consistent across inputs, reducing randomness in relation extraction without strictly enforcing identical wording [29]. We did not set the *top_k* hyperparameter because the LLM should consider all possible completions rather than restrict itself to a limited number of high-probability outputs [30].

After validation, all extracted interactions are post-processed using an automated multi-step rule-based refinement phase to ensure every entity and interaction is expressed using CTD identifiers (S3).

Prompt engineering for this study involved several rounds of iterative refinement to develop the current prompt design. Filtering rules and prompt engineering were initially developed using a set of preliminary articles (PMIDs: 23724058, 24169358, 24664296, 25351418, 25923732, 28285367, 28593498, and 21212295), which are excluded from the final evaluation of 50 articles. Given that the task is complex and the prompts are detailed, we address the ‘‘Lost in the Middle’’ phenomenon, spurious associations, hallucinations, and LLM deterministic traits, further described in S6. We also made key design choices by treating the LLM like a specialized agent, using structured formatting, one- and few-shot learning, and defining clear and concise prompts (S4). Prompt examples are detailed in S9 and S10.

For each extracted interaction, the LLM response includes an explanation, chemical name, chemical ID, interaction, gene name, gene ID, gene symbol, and source text, depending on the data component. Table 2 details input data, output data, and prompt type for each data component for the extraction and validation steps of Clover. The *Avg Tokens* column in Table 2 shows the average tokens used for each prompt. An LLM token for text is a subunit of text, word, or character, and for images, it is a representation of the image using binary data. Images are encoded using base64 encoding, which converts the binary image file into an ASCII string format. A PDF has more tokens because the text and images get compiled into one LLM call. PubTator data is only used to split the PMC articles into paragraphs (i.e., the count value for TEXT, which is 981) and to identify chemicals and genes in the table captions and text.

CTD stores the following information for each interaction: chemical name, chemical ID, CASRN, gene symbol, gene ID, gene forms, organism, organism ID, interaction (a textual representation of the interaction), interaction actions (a list of interaction actions), and PubMed IDs. We do not extract species or gene forms because the LLM often hallucinates the species and/or gene form even when one is absent in the data component during our early experiments.

## Results

First, we compare Clover against a minimal zero-shot whole-PDF baseline to assess differences in overall extraction performance. Then we assess the effect of extraction, validation, and refinement on precision and recall across all models and data components. Finally, we examine model-level differences, relationship distribution, section-type performance, interaction distribution, and entity-level errors to understand where Clover succeeds and fails. In this evaluation, the minimal whole-PDF baseline achieved lower CTD agreement than the evaluated Clover configurations. The metrics are all computationally calculated; additional results are included in S6.

We evaluate Clover against CTD-curated single-step interactions to assess how closely LLM outputs align with existing curation decisions. Therefore, precision and recall are calculated based on exact matches with CTD (PMID, chemical ID, gene symbol, and interaction prefix and type) and should be interpreted as measures of CTD agreement rather than comprehensive biological correctness.

### Effect of Pipeline Stages on Precision and Recall

Fig. 2 is a scatter plot of precision and recall for each standardized and filtered model, with the baseline, which is also standardized and filtered. The Pareto frontier consists of configurations that are not dominated by another configuration in both precision and recall. CL-E (Claude standalone extraction) and CL-V/CL-E (Claude validating Claude extraction output) are the two Pareto-optimal configurations. CL-E achieves the highest precision (61%) with a recall of 71.8%, whereas CL-V/CL-E achieves slightly higher recall at 74.4% with a lower precision of 49%. Given the marginal recall improvement relative to the larger precision decrease, CL-E provides a better balance between precision and recall, which is reflected in its F-1 score (66% for CL-E versus 59% for CL-V/CL-E).

**Fig 2.**
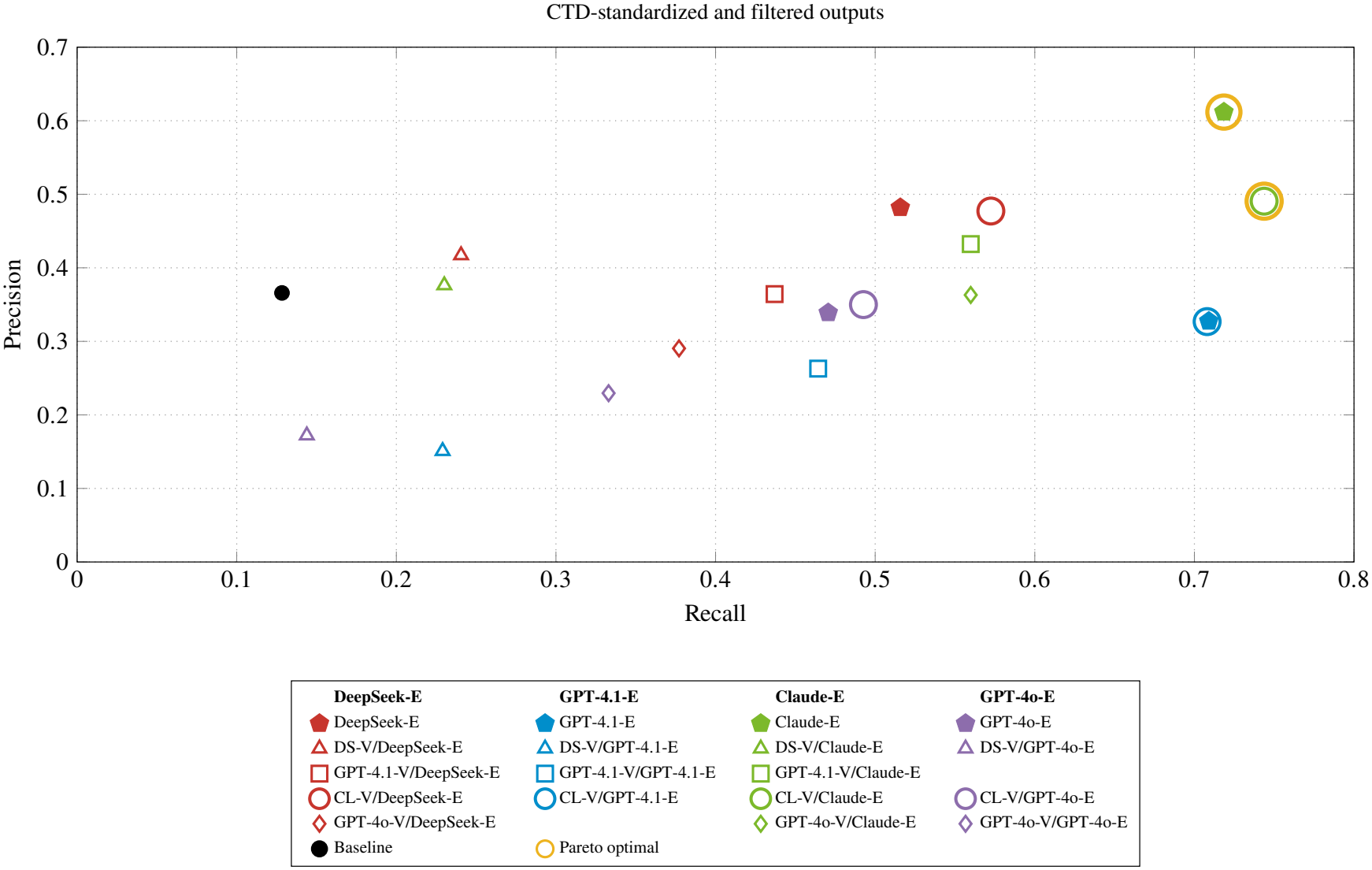
Precision and Recall Scatterplot at CTD-standardized & Filtered stages. This figure shows the precision and recall across all 18 Clover model configurations and the naive baseline, evaluated against CTD-curated chemical-gene interactions for the 50 articles in the evaluation set. The suffixes -E and -V indicate extraction and validation, respectively. Claude is abbreviated to CL and DeepSeek to DS. Extraction-only configurations (e.g., CL-E) and chained configurations (e.g., DS-V/CL-E) undergo both of the refinement steps (standardization and filtering). CL-E and CL-V/CL-E form the Pareto frontier. Claude-Sonnet-4.5 (CL-E) achieves 72% recall and 61% precision. CL-V/CL-E also achieves 74% recall and 49% precision.

Many Clover configurations perform better than the baseline (13% recall and 36% precision) as shown in Fig. 2.

Because the baseline differs in several aspects, including input segmentation, entity annotations, prompt structure, and token length, the observed performance differences cannot be attributed to any single component in our approach. The baseline shows that whole-PDF prompting with minimal guidance is insufficient for CTD-aligned RE.

Fig. 3 shows precision and recall at three checkpoints for all 18 model configurations (with duplicates removed): raw extraction of single-model and validation configurations, after CTD-standardization, and after filtering.

**Fig 3.**
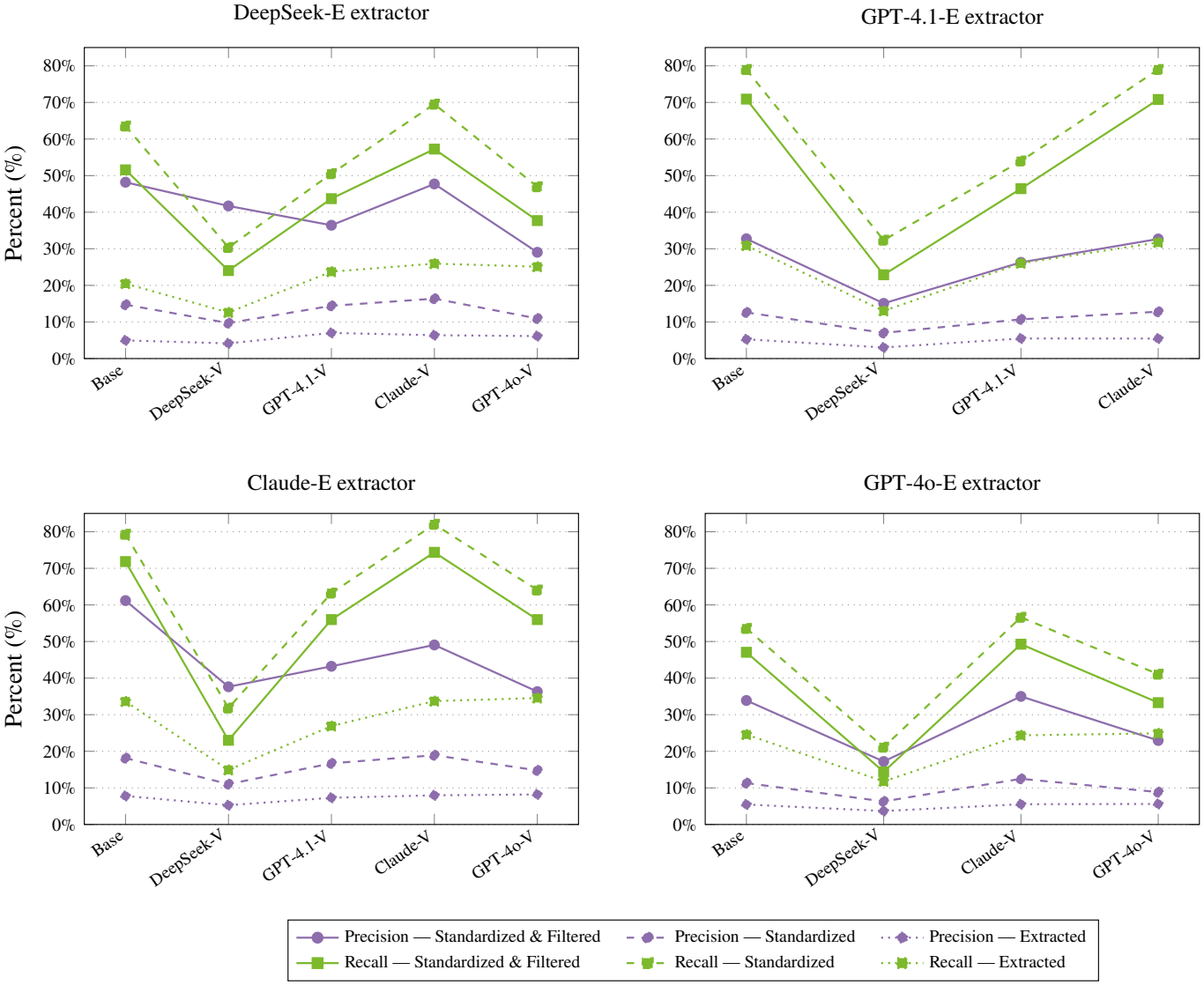
Precision and Recall Line Chart at Extracted, Standardized, and Standardized & Filtered Stages. This figure is a line chart showing precision and recall at three pipeline checkpoints for all single-model and chained configurations with duplicates removed. Each panel represents the extractor model along with the corresponding validator model comparisons. Refinement consistently boosts recall, and filtering substantially increases precision at a modest cost to recall.

#### Single Model Extraction

Across all models, at the extraction stage (dotted line in Fig. 3), both recall and precision are low. The LLMs produce a larger number of candidate interactions, many of which are inconsistent with CTD, unsupported by the data component, missing or invalid entities, and/or assigned the wrong interaction type or prefix, highlighting the improvements observed after postprocessing.

Once the interactions are CTD-standardized (dashed line in Fig. 3), recall is highest across all configurations (79% for GPT-4.1 and Claude, 82% for CL-V/CL-E), and precision is consistently low. Filtering then removes rows citing prior work, describing null or negative findings, entities that are not CTD-standardizable, etc., as shown in Fig. 1. After filtering (solid line in Fig. 3), precision increases substantially and recall declines modestly. Refinement serves as a recall-boosting step, and filtering functions as a high-precision gate.

Claude-Sonnet-4.5 (CL-E) performs the best with 72% recall and 61% precision, making it one of the two Pareto optimal configurations (Fig. 2). The second Pareto optimal configuration is CL-V/CL-E, where Claude self-validates, with 74% recall and 49% precision. GPT-4.1 (GPT-E) is the third strongest, with 71% recall and 33% precision. GPT-4o (GPT-4o-E) has 47% recall and 34% precision, and DeepSeek-V4.1-Flash (DS-E) has 52% recall and 48% precision. DeepSeek and GPT-4o tend to over-generate interactions.

Although we cannot fully attribute the performance gap between Claude and other LLMs, several observations are consistent with Claude’s advantage. First, Claude produces consistent CSV formatting, reducing errors downstream.

Second, the model is more accurate in prefix assignment, setting a prefix to increase or decrease only when the source directly supports it (Fig. 4c in S6). Third, Claude exhibits higher instruction-following fidelity than other models.

**Fig 4.**
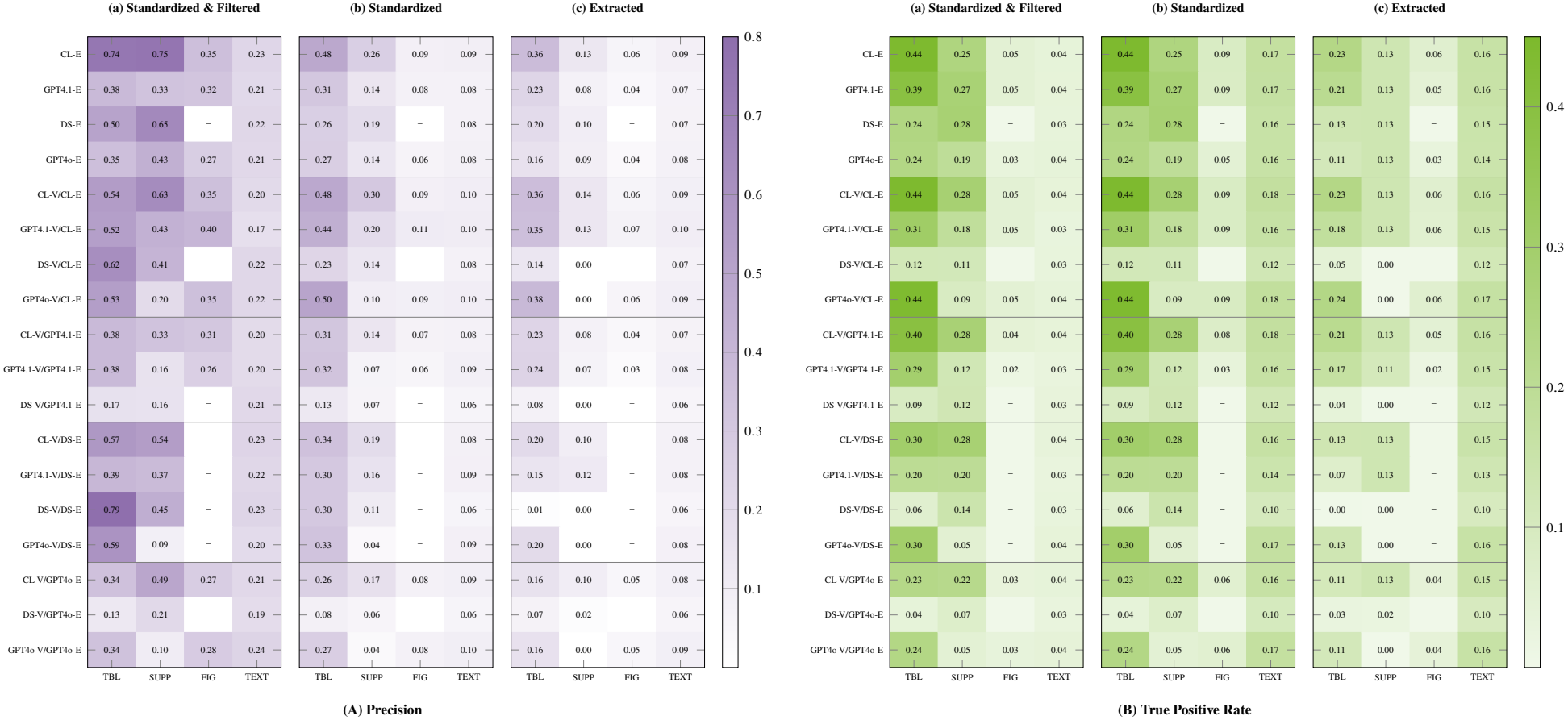
Precision and True Positive Rate. Precision (Fig. 4a) and True Positive Rate (Fig. 4b) matrices broken down by data component (TBL = table, SUP = supplement, FIG = figure, TXT = text) and pipeline stage for all 18 model configurations (for chained configurations, -E = extraction LLM, -V = validation LLM). Structured components (TBL, SUP) produce the highest precision. Text benefits most from filtering. Panel (c) shows each component’s individual contribution before deduplication. **Fig. 4a. Precision matrices by data component and pipeline stage.** **Fig. 4b. True positive rate matrices by data component and pipeline stage.**

The CTD reference contains 952 single-step relationships spanning 17 different interactions, of which 903 (95%) involve expression changes. This imbalance is natural to all of CTD data, where 85% of all single-step relationships are expression-related. Claude matched 662 expression-related relationships and 22 other interaction types. A detailed breakdown of CTD-curated interactions, Claude predictions, and class-level precision and recall is provided in Table 3 in S6 and Fig. 5 in S6.

#### Validation

When a validation model disagrees, it updates the entity or interaction and reflects those changes in the reasoning output column. The chained settings are labeled as Validator(-V)/Extractor(-E) (for example, if DeepSeek is the validator LLM and Claude is the extractor LLM, the label would be DS-V/CL-E).

Validation produces mixed effects on precision and recall depending on the extractor-validator combination. Most chained configurations do not improve performance when compared to their corresponding extraction-only models. However, some combinations provide measurable benefits. For instance, CL-V/DS-E improves recall by 5.7% and reduces precision by a fraction (0.5%), and CL-V/GPT-4o-E improves precision slightly by 1% and recall by 2% relative to GPT-4o-E. Nevertheless, no chained configuration passes Claude-Sonnet-4.5 in precision and recall.

For same-model validation (e.g., CL-V/CL-E), performance decreased for all four LLMs on precision and recall except for CL-E, where recall slightly increased. Specifically, DeepSeek showed the highest recall decrease of 27.5%, removing 316 true positives and gaining only 54. Similarly, GPT-4.1 removed 235 true positives and gained only two. In contrast, Claude removed zero true positives and gained 24, increasing recall by 2.52%. However, Claude gained 308 false positives and removed only seven, resulting in the largest precision decrease of 12.12% (Table 2 in S6).

When a validator is uncertain, it defaults to rejection rather than acceptance, resulting in a net loss.

DeepSeek-V4.1-Flash, as a validator, is a particularly pronounced case of this failure as observed for DS-V/GPT-E, DS-V/GPT-4o-E, and DS-V/CL-E. DeepSeek tends to apply stricter and more literal evidence requirements than other models, frequently flagging interactions as insufficiently supported or changing interaction prefixes.

Chained validation is hypothesized to enhance reliability by filtering erroneous or ambiguous extractions. However, our experiments reveal that only three validation configurations improve precision and/or recall relative to their corresponding extractors. These improvements are model-dependent and do not consistently justify the additional validation step in this BioRE context. The strongest standalone model, Claude, outperforms all evaluated chained configurations.

### Performance Comparison by Data Component

Fig. 4a and Fig. 4b present precision and true positive matrices for Standardized & Filtered data (panel (a)), Standardized data (panel (b)), and Extracted data (panel (c)). Panels (a) and (b) have duplicates removed globally, prioritizing retention of duplicate interactions in structured sources (e.g., tables and supplements) over figures and text. Panel (c) differs because duplicates are removed per section (i.e., an interaction may exist in both table and text), showing where the interactions are extracted from in the PMC article.

#### Structured Data Components

Supplements achieve higher precision compared to other data components across all models since LLMs can parse them easily. Tables have the highest true positive rate across all models and pipeline stages, and moderate precision. A limitation of structured data is the lack of contextual information, which prevents the LLM from identifying where an interaction originates within a table or supplement. Therefore, it is difficult to filter out unsupported interactions. As a result, the number of incorrect tabular entries is high (Fig. 4a in S6).

#### Unstructured Data Components

Figure data achieve precision ranging from 0.26 to 0.41 (Fig. 4a). DeepSeek does not support images. Figures are processed without NER annotations because PubTator does not support images (Table 2). Image content often contains bar graphs (e.g., western blots) or pathway diagrams, which require the LLM to interpret graphical conventions that are often less standardized. Despite these challenges, figures contribute to some correct extractions for well-labeled diagrams where chemical and gene names appear in the caption or the image. Text sections have the most dramatic post-filter precision increase (Panels (b) and (a) of Fig. 4a) due to removing frequent references to prior work and/or common knowledge, and descriptions of non-significant experimental results.

#### Interactions identified beyond CTD

Clover may extract interactions that have not been recorded by CTD. We performed a manual analysis to assess how often Clover identifies potentially valid interactions absent from CTD. A manual review was conducted on the text data component for CL-E. We randomly selected 200 chemical–gene relationships extracted from text by Claude-E (CL-E) for manual review. We then examined the corresponding predictions from all four standalone extraction models to identify potentially valid interactions absent from CTD and found that CL-E has four, GPT-E and GPT-4o-E have three each, and DS-E has one such interaction. The manual review was conducted by Allan Peter Davis, CTD Biocuration Project Manager and Co-Principal Investigator, who has over 20 years of CTD biocuration experience. The analysis examined whether CTD-unmatched predictions were supported by the source text and consistent with CTD curation criteria. Because the review was conducted by a single expert, no inter-annotator disagreement resolution was necessary.

Unmatched predictions from tables, figures, and supplements were not systematically reviewed. The reported precision and recall in this work measure agreement with CTD curation rather than the complete scientific validity of extracted interactions. A human curator reviewing the LLM output may add additional interactions to CTD rather than discard them, especially for table and figure sections, which were not manually evaluated for additional interactions. Additionally, a previous CTD inter-biocurator study reported an average precision of 91% and recall of 71% against an expert-adjudicated reference, highlighting variability in manual curation coverage [3].

### Qualitative Case Studies: Mismatches Between Clover and CTD

Across the examples, there are three distinct types of mismatches between Clover and CTD-curated data. First is an *interaction type* mismatch where the chemical-gene pair and directionality are correct, but the model selects a different interaction because the source text uses informal language that steers the model towards a generic label (Example 1). Alternatively, the article’s framing describes the finding in terms of drug resistance rather than direct molecular regulation that CTD annotates at a different level of specificity compared to an LLM (Example 2).

Second are *entity resolution failures*, where the model identifies the correct entity, but the refinement step resolves to an incorrect entity—either to a close family member (Example 3), an unrelated record (Example 4), or CTD biocurators assigned another entity of the same family (Example 5). Third are *curation scope gaps*, where a model extracts the correct chemical-gene pair and interaction, but it is not curated by CTD because the chemical is not a primary focus of the PMC article (Example 6) or because CTD curates statistically significant data (Example 7).

The first and second categories present engineering problems. The third suggests that precision against CTD may be a lower bound on Clover’s true utility, as a non-trivial fraction of false positives correspond to interactions that CTD has not yet curated. Since CTD curates interactions according to its policies, some unmatched predictions may be scientifically supported but outside CTD’s curation scope. Systematic expert review for all model extractions is needed to distinguish these from genuine extraction errors. The following examples are from CL-E standardized and filtered output, illustrating model failures across all data components. The **C****lover****Output [Chemical–Interaction–Gene]** is the interaction from Claude output (CL-E).

Clover Example Output 1: Correct Entities, Wrong Relationship Type

**PMID:** 21294877

**Data Component:** ABSTRACT (TEXT)

**C****lover** **Output [Chemical–Interaction–Gene]:**

Lipopolysaccharides– increases activity– TAC1

**CTD-curated Interaction:** increases expression

**LLM-cited Evidence:** *‘‘In contrast, induction of the TAC1 promoter by LPS was not dependent on ECR2 and did not occur in large diameter neurones*.*’’*

**Analysis:** The chemical-gene pair and direction are correct, but the interaction type differs. Claude interprets induction of the TAC1 promoter as increased activity, whereas CTD records increased expression of TAC1. This illustrates how promoter activity and gene expression can be interpreted differently during relation extraction.

Clover Example Output 2: Correct Entities, Different Relationship

**PMID:** 23349674

**Data Component:** TABLE

**C****lover** **Output [Chemical–Interaction–Gene]:**

Arsenic– affects expression– CAPN10

**CTD-curated Interaction:** affects response to substance

**Table caption:** *‘‘Univariate and multivariate association between alleles for SNPs-43 and -44 in CAPN-10 and Log-beta cell function (%) in non-diabetic and diabetic subjects*.*’’*

**Analysis:** Claude identifies the correct chemical-gene pair but assigns an expression-related interaction. CTD instead curates the relationship as an effect of CAPN10 genetic variation on susceptibility to arsenic. Both relationships are technically correct; however, the CTD-curated statement reflects additional domain-based expertise to achieve greater specificity.

Clover Example Output 3: Wrong Gene Family Member Resolved

**PMID:** 18299717

**Data Component:** TABLE

**C****lover** **Output [Chemical–Interaction–Gene]:**

Rebamipide–increases expression–AKR1C3

**CTD-curated Gene:** AKR1C14

**Table caption:** *‘‘Genes up-regulated at least 1*.*5-fold in rebamipide treatment in rats’’*

**Analysis:** The table indicates that rebamipide increases expression of *‘‘3-alpha-hydroxysteroid dehydrogenase’’*. Since no NER exists inside a table, the gene-matching code outputs AKR1C3. Without a more specific identifier in the LLM output—and compounded by the fact that Clover does not extract species, which may help narrow to the correct family member—the refinement step selects the wrong isoform.

Clover Example Output 4: Missing Gene from Figure

**PMID:** 24413757

**Data Component:** FIGURE

**C****lover** **Output [Chemical–Interaction–Gene]:** Mipafox–decreases activity–NTE

**CTD-curated Gene:** PNPLA6

**Figure caption:** *Changes in NTE activity of the NT2 cells exposed to paraoxon or mipafox during the neurodifferen-tiation process*.

**Analysis:** CTD confirms the correct entry is mipafox–decreases activity–PNPLA6, where PNPLA6 encodes NTE. The figure caption and image refer to ‘‘NTE’’ and not PNPLA6. Therefore, the model relies on the figure caption and image data provided, resolving to NTE.

Clover Example Output 5: Gene Symbol Resolution

**PMID:** 22747577

**Data Component:** SUPPLEMENT

**C****lover** **Output [Chemical–Interaction–Gene]:**

Masitinib–decreases expression–UQCC

**CTD-curated Gene:** UQCC1

**Supplement caption:** *‘‘Genes with a time-dependent, continuous increase or decrease in mRNA expression level during masitinib treatment*.*’’*

**Analysis:** The discrepancy arises because the supplemental data lists the gene as UQCC, which the system extracted correctly, but reflects an incorrect use of gene terminology in the manuscript, which CTD curators can rectify.

Clover Example Output 6: Simplified Multistep Extraction

**PMID:** 17524151

**Data Component:** TEXT

**C****lover** **Output [Chemical–Interaction–Gene]:**

Roxithromycin–decreases expression–CXCL8

**LLM-cited Evidence:** *‘‘roxithromycin inhibited the SM-stimulated overproduction of the proinflammatory cy-tokines…IL-8…’’*

**Analysis:** The sentence clearly shows roxithromycin decreases IL-8 (CXCL8) matching *decreases expression*. The mismatch reflects the more detailed scope of manual curation, as the relationship between Roxithromycin and CXCL8 is multistep in CTD; multistep relationships are out of scope for this work.

Clover Example Output 7: Rare Relationship Type

**PMID:** 25859307

**Data Component:** FIGURE

**C****lover** **Output [Chemical–Interaction–Gene]:**

Ethanol– increases cleavage– CASP3

**CTD-curated Interaction:** increases cleavage

**Figure caption:** *‘‘Effect of NSO (0*.*5 ml/kg) and ethanol on the protein level of caspase-3 (cleaved caspase-3) in the rat liver tissue*.*’’*

**Analysis:** Claude correctly identifies the increased cleavage of CASP3 following ethanol exposure, matching the CTD-curated relationship. Unlike expression-related interactions, cleavage represents a less frequently observed relationship type in our evaluation. This example shows that the model can capture less common relationship types, although in a less nuanced manner.

## Discussion

Clover’s results demonstrate the potential of using LLMs for CTD-aligned chemical-gene interaction extraction from full-text PMC articles. Benchmarking four state-of-the-art general-purpose LLMs (GPT-4.1, GPT-4o, Claude-Sonnet-4.5, and DeepSeek-V4.1-Flash), we show the evaluated configurations outperform the minimal whole-PDF baseline (13% recall, 36% precision), with Claude-Sonnet-4.5 achieving 72% recall and 61% precision.

However, because the baseline differs in multiple aspects, these results do not establish the independent contribution of structured prompting, PubTator NER, or section-level processing.

Validation improves performance for only three extractor-validator combinations. These benefits are model-dependent and are not observed consistently across configurations. In real-world deployment, chaining also increases API costs and processing latency for low returns. The Refinement stage, consisting of CTD-standardization and Filtering, substantially improved the measured performance.

We accessed DeepSeek-V4.1-Flash through a hosted API in our experiments. However, it is publicly available for local deployment. Since DeepSeek-V4.1-Flash has 671 billion total parameters, it requires substantial GPU memory and computational resources. Therefore, local hosting is impractical under our current resource constraints. Future work could evaluate smaller, locally deployable open-weight models to determine whether comparable Clover performance can be achieved.

Qualitative case study analysis reveals three recurring mismatch categories—(i) when informal language steers the model towards a broader label or the model annotates an interaction at a different biological level compared to a curator, (ii) when the gene-matching algorithm resolves to an incorrect gene, and (iii) when models sometimes extract interactions differently and typically more simply than curators.

There are several important strengths to our study. This is the first benchmark for CTD-aligned full-text chemical-gene interaction extractions. Although some chained configurations improved upon their corresponding extraction-only models, Claude alone achieved the strongest overall performance. Refinement improved performance across evaluated pipeline stages. We also found that implementing rule-based refinement post-extraction substantially increased performance metrics.

As well, there are some important limitations to consider.

1. *Incomplete Article Context* CTD curators extract interactions from the entirety of an article, including figures, tables, and text. In contrast, Clover has visibility into individual sections of the article at a time. Therefore, any interaction inferred from multiple passages across the paper is inherently invisible to the LLM.
2. *Small and Restricted Corpus* We tested Clover against a small corpus of 50 scientific articles, four of which contained a CSV-based supplemental table; this limited sample size is not necessarily representative of the 150,000 manually curated articles currently in CTD. The imbalance of expression-related interactions limits conclusions about rare interaction classes and may affect the stability of model rankings on different datasets.
3. *Incomplete Interaction Scope* Our study addressed only a fraction of the scope of content that is manually captured by CTD biocurators. Specifically, Clover was restricted to finding tripartite interaction components (chemical-relationship-gene), whereas CTD biocurators also identify additional qualifiers such as the gene-form being affected (e.g., mRNA, protein, DNA, promoter, etc.), how the interaction was measured experimentally (e.g., in vivo or in vitro), and the organism (species) in which the interaction occurred; all of these additional data fields have associated controlled vocabularies. More importantly, CTD biocurators typically curate complex interactions that involve more than a single chemical, gene, and interaction type within the relationship [31], such as: ‘‘Chemical 2 promotes [Chemical 1 inhibits [Gene 1 protein increases the expression of Gene 2 mRNA]],’’ which involves two chemicals, two genes, two gene-forms, three interaction types, and one organism.
4. *Incomplete Reference and Adjudication:* CTD-curated interactions are not an exhaustive reference for all biologically valid relationships. Our evaluation measures exact CTD agreement, and manual examination of additional relationships was limited to text-derived predictions. Therefore, the biological validity of all predictions not matching CTD remains unverified.
5. *Limited Source Coverage* Clover is currently limited to papers available in PubMed Central (roughly one-third of CTD-curated papers), because PubTator relies on the PMC Open Access Subset for full-text processing. CTD biocurators also manually extract data for other modules—including chemical-phenotype, gene-disease, and the 33-component exposure module—none of which are addressed here [32].

### Potential CTD Integration

Clover could help CTD curators by suggesting candidate chemical-gene interactions tagged at the appropriate source (figure, text, table, abstract) in full-text articles before manual curation. These suggested interactions could then be reviewed by curators to determine if they should be added to CTD. The curation process would still be performed by curators who would be responsible for making sure that the interaction is present in the text and meets CTD’s standards for curation, including considering factors beyond what Clover currently extracts and determines as a candidate. Clover could help curators prioritize LLM candidate interactions before manually finding interactions mentioned in the text. However, the impact of Clover would need to be tested within CTD to see whether the time or effort required from curators is reduced.

Considerably more work is needed to advance LLM-based tools to approximate the full capacity and extent of manual curation at CTD. Finally, it should be noted that the effectiveness of LLMs going forward will be constrained by the lack of standardization of key elements across the publishing industry. For example, entities like chemical, gene, and disease term names are often left solely to authors’ discretion; similarly, the formats of the various components that comprise machine-readable full text and supplements vary by publisher. Standardizing these types of components and entities would greatly facilitate LLM processing.

## Conclusion

This work introduces Clover, a training-free human-review pipeline that harnesses LLMs to extract CTD-aligned chemical-gene interactions from full-text PMC articles. Clover uses CTD-defined interaction terminology for full-text chemical-gene relation extraction, integrating PubTator-grounded named entity recognition, structured one-shot and few-shot prompting, and refinement.

Using four state-of-the-art LLMs (GPT-4.1, GPT-4o, Claude-Sonnet-4.5, and DeepSeek-V4.1-Flash) in single-model and chained-model configurations, we find that Claude-Sonnet-4.5 alone achieves the highest performance (61% precision and 72% recall). Although CL-V/CL-E achieves slightly higher recall, this improvement comes with a substantial precision decrease. Chaining models produces mixed performance changes, with improvements observed for some extractor-validator combinations. Therefore, an additional validation stage is not necessary to achieve the strongest performance among the evaluated configurations. Instead, Claude-Sonnet-4.5 alone performs the best and reduces API costs and latency. Potentially, Clover could assist CTD curators by providing candidate interactions for review, although its impact on curator workload has not yet been evaluated. Practical implementation details are outlined in S6.

Clover extends what is already possible with PubTator alone. PubTator performs NER for biomedical entities; however, it does not extract the interactions between them at the specificity required for CTD curation. Clover builds on this by performing relation extraction from full-text articles. The refinement steps could potentially reduce the burden on curators by presenting candidate interactions for review, accelerating the rate at which curated knowledge enters databases.

Additionally, Clover has the potential to accelerate CTD updates. Clover is unsupervised and thereby reduces dependency on large annotated datasets for training and may be a practical solution for large-scale data curation efforts. The present work opens up future directions to enhance CTD alignment.

We propose several enhancements to this reported work, including specialized LLM Chains, improved context, fine-tuning, and multistep interaction extraction.

1. *Controlled Ablations* Future work could isolate the contributions of structured prompting, PubTator NER, section splitting, and refinement through controlled ablations.
2. *Specialized LLM Chains* Instead of using a single LLM for extraction, we can employ multiple LLMs (or multiple prompt stages) focusing on different components of the interaction, e.g., entity pairs, interaction prefix, and interaction type.
3. *Improved Context* To address the limitation of limited context, we can preprocess text by merging relevant passages to provide additional information for the LLM.
4. *Fine-Tuning LLMs* Instead of relying solely on prompt-based approaches, future work can involve fine-tuning LLMs using CTD data to use domain-specific knowledge.
5. *Multistep Interaction Extraction* Clover currently supports single-step interactions. Multistep interactions may be extracted by first identifying candidate entity chains within a passage and then prompting the LLM to characterize each link in the chain independently before assembling the full multi-step interaction.
6. *Curator-C**lover* *Comparison* Future work could evaluate Clover alongside routine CTD curation, allowing direct comparisons between model-generated interactions and curator decisions across a larger and more diverse set of articles. This evaluation would also help identify reliable article and evidence types to quantify whether Clover reduces the time and effort required for manual curation.
7. *Experimental Context Extraction* Future work could extend Clover to extract experimental metadata, including tissue, organ, cell type, organism, in vivo or in vitro setting, exposure dose, and duration. This information would provide additional biological context for interpreting chemical-gene relationships and for use in downstream analyses.

With these enhancements, we could potentially implement Clover as a semi-automated assistant for CTD curation, helping scale biomedical knowledge to support drug discovery, clinical research, and toxicogenomics.

## Supporting information

System prompt for figure extraction.

User prompt for figure extraction.

System prompt for table extraction

User prompt for table extraction

System prompt for supplement extraction

User prompt for supplement extraction

System prompt for text extraction

User prompt for text extraction

System prompt for figure validation

User prompt for figure validation

System prompt for table validation.

User prompt for table validation.

System prompt for supplement validation

User prompt for supplement validation

System prompt for text validation.

User prompt for text validation.

## Acknowledgments

The authors thank Sakib Abrar for his helpful comments and discussions.

Generative AI Disclosure Statement: AI tools were used to assist with code development, debugging, and figure preparation. Generative AI was not used to develop the experimental prompts. For figures, the authors used AI assistance to update existing figure templates in LATEX using author-generated data. All AI-assisted content was reviewed and verified by the authors, who take full responsibility for the final manuscript.

## Supporting information

**S1 Appendix. Supplementary methods and results for C****lover**. Additional details on the naive baseline, rule-based refinement, prompt design, article-level evaluation, model comparisons, gene normalization, filtering-loss analysis, implementation, and extraction and validation prompts.

**S2 Fig. 1** List of interactions using CTD classification scheme.

**S4 Fig. 2** Negative and null filtering patterns used in the Refinement stage to remove interactions where the source text indicates no effect, failed outcomes, undetectable measurements, contradictory associations, or unconfirmed findings. These patterns do not automatically exclude decreases in measurable gene activity or expression.

**S5 Fig. 3** Broad and general statement filtering patterns used in the Refinement stage to remove interactions where the source text reflects common knowledge, hedged associations, or references to prior literature rather than direct experimental evidence.

**S6 Fig. 4a Standardized & Filtered stacked bar chart**. Standardized & Filtered stacked bar chart showing the count of correct (solid), partial match (hatched), and incorrect (transparent) interactions per data component (TABLE, SUPPLEMENT, FIGURE, TEXT) after CTD-standardization and filtering steps.

**S6 Fig. 4b Standardized-only stacked bar chart**. Standardized-only stacked bar chart (duplicates removed with structured-data priority, no filtering) showing the substantially larger candidate set produced before filtering, illustrating how filtering reduces the candidate interaction set.

**S6 Fig. 4c Invalid entity and interaction error counts**. Invalid entity and interaction error counts per configuration at the Standardized & Filtered stage, decomposed into five error classes: invalid gene symbols, invalid chemical IDs, wrong prefix only, wrong interaction type only, and wrong prefix and type combined. Invalid gene symbols are the most prevalent error class, reflecting the difficulty of resolving gene identifiers from informal or ambiguous mentions in biomedical text. Error categories overlap; therefore, the sum of errors cannot be used to obtain the total number of incorrect predictions. CL-E produces fewer prefix-only errors than other models. Validation does not reduce interaction errors, supporting the finding that LLM chaining is not an effective strategy for this task.

**S6 Fig. 5 Relationship Type Distribution per Model**. Stacked bar chart comparing the distribution of interaction types across all 18 Clover model configurations and CTD-curated relationships for the 50 PMC articles, shown at the Standardized & Filtered stage. Relationship types are color-coded, with the most frequent CTD-curated types highlighted. CTD-curated relationship distribution is heavily concentrated in expression-related interactions, a pattern broadly reflected across model outputs. Model distributions match CTD exact-matched chemical, gene, and interaction relationships.

**S8 Fig. 6 Naive baseline prompt**. Naive baseline prompt applied directly to full-text PDFs. Unlike Clover, this prompt provides no PubTator NER, no section splitting, no entity grounding, and no examples.

**S9 Fig. 7** System prompt for figure extraction.

**S9 Fig. 8** User prompt for figure extraction.

**S9 Fig. 9** System prompt for table extraction.

**S9 Fig. 10** User prompt for table extraction. LLMs perform better with raw XML tables because tables are usually complex (may include multiple headers), and these do not parse properly.

**S9 Fig. 11** System prompt for supplement extraction.

**S9 Fig. 12** User prompt for supplement extraction.

**S9 Fig. 13** System prompt for text extraction.

**S9 Fig. 14** User prompt for text extraction.

**S10 Fig. 15** System prompt for figure validation.

**S10 Fig. 16** Structured validation prompt for figure data using one-shot learning.

**S10 Fig. 17** System prompt for table validation.

**S10 Fig. 18** Structured validation prompt for table data using one-shot learning.

**S10 Fig. 19** System prompt for supplement validation.

**S10 Fig. 20** Structured validation prompt for CSV supplement data using one-shot learning.

**S10 Fig. 21** System prompt for text validation.

**S10 Fig. 22** Structured validation prompt for text data using one-shot learning.

## Notes

https://github.com/sanjucheerla/CLOVER/

### Competing Interest Statement

The authors have declared no competing interest.

### Funding

- National Institute of Environmental Health Sciences: U24ES033155

## References

1. Davis AP, Wiegers TC, Sciaky D, Barkalow F, Strong M, Wyatt B, et al. Comparative Toxicogenomics Database’s 20th Anniversary: Update 2025. Nucleic Acids Research. 2025;53(D1):D1328–34.

2. Nachtegael C, De Stefani J, Lenaerts T. A Study of Deep Active Learning Methods to Reduce Labelling Efforts in Biomedical Relation Extraction. PLOS ONE. 2023 12;18(12):1–23. Available from: https://doi.org/10.1371/journal.pone.0292356. doi:10.1371/journal.pone.0292356.

3. Wiegers TC, Davis AP, Cohen KB, Hirschman L, Mattingly CJ. Text Mining and Manual Curation of Chemical-Gene-Disease Networks for the Comparative Toxicogenomics Database (CTD). BMC Bioinformatics. 2009;10:326. doi:10.1186/1471-2105-10-326.

4. Luo R, Sun L, Xia Y, Qin T, Zhang S, Poon H, et al. BioGPT: Generative Pre-Trained Transformer for Biomedical Text Generation and Mining. Briefings in Bioinformatics. 2022 09;23(6):bbac409. Available from: https://doi.org/10.1093/bib/bbac409. arXiv:https://academic.oup.com/bib/article-pdf/23/6/bbac409/47144271/bbac409.pdf. doi:10.1093/bib/bbac409.

5. Zhao X, Deng Y, Yang M, Wang L, Zhang R, Cheng H, et al. A Comprehensive Survey on Relation Extraction: Recent Advances and New Frontiers. ACM Computing Surveys. 2024 jul;56(11). Available from: https://doi.org/10.1145/3674501. doi:10.1145/3674501.

6. Ono T, Hishigaki H, Tanigami A, Takagi T. Automated Extraction of Information on Protein–Protein Interactions from the Biological Literature. Bioinformatics. 2001;17(2):155–61.

7. Huang MS, Han JC, Lin PY, You YT, Tsai RTH, Hsu WL. Surveying Biomedical Relation Extraction: A Critical Examination of Current Datasets and the Proposal of a New Resource. Briefings in Bioinformatics. 2024 04;25(3):bbae132. Available from: https://doi.org/10.1093/bib/bbae132. arXiv:https://academic.oup.com/bib/article-pdf/25/3/bbae132/57221564/bbae132.pdf. doi:10.1093/bib/bbae132.

8. Devlin J, Chang M, Lee K, Toutanova K. BERT: Pre-Training of Deep Bidirectional Transformers for Language Understanding. In: Burstein J, Doran C, Solorio T, editors. Proceedings of the 2019 Conference of the North American Chapter of the Association for Computational Linguistics: Human Language Technologies (NAACL-HLT 2019). Association for Computational Linguistics; 2019. p. 4171–86. Available from: https://doi.org/10.18653/v1/n19-1423. doi:10.18653/V1/N19-1423.

9. Lee J, Yoon W, Kim S, Kim D, Kim S, So CH, et al. BioBERT: A Pre-Trained Biomedical Language Representation Model for Biomedical Text Mining. Bioinformatics. 2019;abs/1901.08746. Available from: http://arxiv.org/abs/1901.08746. arXiv:1901.08746.

10. Jahan I, Laskar MTR, Peng C, Huang JX. A Comprehensive Evaluation of Large Language Models on Benchmark Biomedical Text Processing Tasks. Computers in Biology and Medicine. 2024;171:108189. doi:10.1016/j.compbiomed.2024.108189.

11. Asada M, Fukuda K. Enhancing Relation Extraction from Biomedical Texts by Large Language Models. In: International Conference on Human-Computer Interaction. Springer; 2024. p. 3–14.

12. Zhou H, Li M, Xiao Y, Yang H, Zhang R. LEAP: LLM Instruction-Example Adaptive Prompting Framework for Biomedical Relation Extraction. Journal of the American Medical Informatics Association. 2024;31(9):2010–8. Available from: https://doi.org/10.1093/jamia/ocae147. doi:10.1093/JAMIA/OCAE147.

13. Zhang D, Mohan S, Torkar M, McCallum A. A Distant Supervision Corpus for Extracting Biomedical Relationships Between Chemicals, Diseases and Genes. In: Calzolari N, Béchet F, Blache P, Choukri K, Cieri C, Declerck T, et al., editors. Proceedings of the Thirteenth Language Resources and Evaluation Conference. Marseille, France: European Language Resources Association; 2022. p. 1073–82. Available from: https://aclanthology.org/2022.lrec-1.116/.

14. Dong X, Zhao D, Meng J, Guo B, Lin H. SyRACT: Zero-Shot Biomedical Document-Level Relation Extraction with Synergistic RAG and CoT. Bioinformatics. 2025;41(7):btaf356. doi:10.1093/bioinformatics/btaf356.

15. Brokman A, Ai X, Jiang Y, Gupta S, Kavuluru R. A Benchmark for End-to-End Zero-Shot Biomedical Relation Extraction with LLMs: Experiments with OpenAI Models. In: Proceedings of the Third Workshop for Artificial Intelligence for Scientific Publications. Association for Computational Linguistics; 2025. p. 44–55. Available from: https://aclanthology.org/2025.wasp-main.6/. doi:10.18653/v1/2025.wasp-main.6.

16. Wei CH, Allot A, Lai PT, Leaman R, Tian S, Luo L, et al. PubTator 3.0: An AI-Powered Literature Resource for Unlocking Biomedical Knowledge. Nucleic Acids Research. 2024 04;52(W1):W540–6. Available from: https://doi.org/10.1093/nar/gkae235. arXiv:https://academic.oup.com/nar/article-pdf/52/W1/W540/58436124/gkae235.pdf. doi:10.1093/nar/gkae235.

17. National Center for Biotechnology Information. PubMed Central; 2024. Accessed: 2025-03-02. https://pmc.ncbi.nlm.nih.gov/.

18. Wadhwa S, Amir S, Wallace B. Revisiting Relation Extraction in the Era of Large Language Models. In: Proceedings of the 61st Annual Meeting of the Association for Computational Linguistics (Volume 1: Long Papers). Toronto, Canada: Association for Computational Linguistics; 2023. p. 15566–89. Available from: https://aclanthology.org/2023.acl-long.868/. doi:10.18653/v1/2023.acl-long.868.

19. Zhang J, Wibert M, Zhou H, Peng X, Chen Q, Keloth VK, et al. A Study of Biomedical Relation Extraction Using GPT Models. AMIA Summits on Translational Science Proceedings. 2024;2024:391.

20. Zhao D, Yang Y, Chen P, Meng J, Sun S, Wang J, et al. Biomedical Document Relation Extraction with Prompt Learning and KNN. Journal of Biomedical Informatics. 2023;145:104459.

21. Choudhury RR, Dey S. GPT-3 Powered Information Extraction for Building Robust Knowledge Bases. arXiv preprint arXiv:240804641. 2024.

22. Chen Q, Hu Y, Peng X, Xie Q, Jin Q, Gilson A, et al. Benchmarking Large Language Models for Biomedical Natural Language Processing Applications and Recommendations. Nature Communications. 2025;16(1):3280.

23. Gill JK, Chetty M, Lim S, Hallinan J. Large Language Model Based Framework for Automated Extraction of Genetic Interactions from Unstructured Data. PLOS ONE. 2024 05;19(5):1–22. Available from: https://doi.org/10.1371/journal.pone.0303231. doi:10.1371/journal.pone.0303231.

24. Laskar MTR, Jahan I, Dolatabadi E, Peng C, Hoque E, Huang JX. Improving Automatic Evaluation of Large Language Models (LLMs) in Biomedical Relation Extraction via LLMs-as-the-Judge. In: Proceedings of the 63rd Annual Meeting of the Association for Computational Linguistics (Volume 1: Long Papers). Association for Computational Linguistics; 2025. p. 25483–97. Available from: https://aclanthology.org/2025.acl-long.1238/. doi:10.18653/v1/2025.acl-long.1238.

25. Dip SA, Mallick D, Shuvo UA, Soumma SB, Rafsani F, Paul BK, et al. Large Language Model Agents for Biological Intelligence Across Genomics, Proteomics, Spatial Biology, and Biomedicine. Briefings in Bioinformatics. 2026;27(2):bbag110. doi:10.1093/bib/bbag110.

26. Zhang Y, Sun R, Chen Y, Pfister T, Zhang R, Arik S. Chain of Agents: Large Language Models Collaborating on Long-Context Tasks. Advances in Neural Information Processing Systems. 2024;37:132208–37.

27. Chen M, Zhang T, Wang S. Prompting Large Language Models to Extract Chemical-Disease Relation Precisely and Comprehensively at the Document Level: An Evaluation Study. PLOS ONE. 2025;20(4):e0320123. Available from: https://doi.org/10.1371/journal.pone.0320123. doi:10.1371/journal.pone.0320123.

28. Grunde-McLaughlin M, Lam MS, Krishna R, Weld DS, Heer J. Designing LLM Chains by Adapting Techniques from Crowdsourcing Workflows. arXiv preprint arXiv:231211681. 2023.

29. Renze M. The Effect of Sampling Temperature on Problem Solving in Large Language Models. In: Findings of the Association for Computational Linguistics: EMNLP 2024. Miami, Florida, USA: Association for Computational Linguistics; 2024. p. 7346–56. Available from: https://aclanthology.org/2024.findings-emnlp.432/. doi:10.18653/v1/2024.findings-emnlp.432.

30. Peeperkorn M, Kouwenhoven T, Brown D, Jordanous A. Is Temperature the Creativity Parameter of Large Language Models? arXiv preprint arXiv:240500492. 2024.

31. Davis AP, Wiegers TC, Rosenstein MC, Murphy CG, Mattingly CJ. The Curation Paradigm and Application Tool Used for Manual Curation of the Scientific Literature at the Comparative Toxicogenomics Database. Database. 2011;2011:bar034. doi:10.1093/database/bar034.

32. Grondin CJ, Davis AP, Wiegers TC, King BL, Wiegers JA, Reif DM, et al. Advancing Exposure Science through Chemical Data Curation and Integration in the Comparative Toxicogenomics Database. Environmental Health Perspectives. 2016;124(10):1592–9. doi:10.1289/EHP174.

