## Supplement 1 for "CLOVER: Chemical-gene LLM benchmark for obtaining valid entity relationships for the Comparative Toxicogenomics Database (CTD)"

### Supporting information

**S1 Appendix.** Supplementary methods and results for CLOVER. Additional details on the naive baseline, rule-based refinement, prompt design, article-level evaluation, model comparisons, gene normalization, filtering-loss analysis, implementation, and extraction and validation prompts.

**S2 Fig. 1** List of interactions using CTD classification scheme.

**CTD-aligned Interaction List**

cotreatment, binding, reaction, activity, localization, expression, abundance, mutagenesis, response to chemical, stability, splicing, folding, transport, uptake, import, secretion, export, metabolic processing, chemical synthesis, degradation, acetylation, acylation, alkylation, amination, carbamoylation, carboxylation, cleavage, ethylation, glycation, glycosylation, N-linked glycosylation, O-linked glycosylation, glucuronidation, hydrolysis, hydroxylation, lipidation, geranoylation, farnesylation, myristoylation, palmitoylation, prenylation, methylation, nitrosation, nucleotidylation, oxidation, phosphorylation, sulfation, reduction, ribosylation, ADP-ribosylation, ubiquitination, sumoylation, glutathionylation, polymerization

**Fig 1.** List of interactions using CTD classification scheme.

**S3 Appendix.** Naive Baseline

Since there is no previous work extracting chemical-gene interactions from full-text articles using the CTD-aligned data classification scheme, we compare our results to a naive baseline. The baseline uses PDF versions of the 50 full-text articles, which were extracted using a web scraper that we built. We uploaded each PDF along with a minimal unstructured prompt explaining that the model must extract interactions between chemicals and genes present in the article.

Unlike Clover, the naive baseline does not use PubTator for NER and does not split the article into sections. We apply the same refinement process used in Clover (described in the Clover Approach section) to compare our results with CTD-curated data. The baseline prompt specifies the output format and the set of valid interaction types, but provides no examples, no entity grounding, and no structural guidance. The baseline design is intentional—reflecting the minimum viable effort a curator might apply when using an LLM for relation extraction. Both approaches undergo the same refinement process, but differences in preprocessing and prompting prevent attribution of performance gains to individual components.

**S4 Appendix.** Rule-Based Refinement

**CTD Standardization:** Refinement consists of CTD standardization and Filtering. In CTD standardization, the entities and interactions are updated to match CTD vocabulary.

**Chemicals:** The process first checks if a Chemical ID is present in the extracted output. If one is found, the chemical name is updated to match the official name recorded in CTD for that ID. Otherwise, the chemical name itself is used to search CTD chemical data. First, we computationally search against known chemical names and then against CTD-Curated and MeSH synonyms, and the correct ID is written back. Any interaction where a valid Chemical ID cannot be resolved is flagged for removal.

**Genes:** A three-step priority chain is applied. First, if a Gene ID is present in the extracted output and can be found directly in CTD, the gene symbol and name are updated to match the CTD record for that ID, since a direct ID match is considered highly reliable. Second, if the Gene ID is not available, the gene symbol is looked up next, and the corresponding ID, symbol, and name are pulled from CTD.

Third, if neither the ID nor the symbol resolves to a match, the gene name is used. We computationally search against both gene names and synonyms in CTD. If the gene name matches multiple CTD gene names and IDs, we choose the CTD gene that has the most alternative IDs as a confidence signal. Alternative IDs in CTD are NCBI gene IDs, each of which corresponds to a specific species.

We choose the gene with the highest count of alternative IDs to maximize species choices since the LLM does not output species. As a further cleaning step for any genes resolved from step three, we strip the dot suffix back to its base symbol and re-resolve against CTD to retrieve the correct Gene ID (e.g. `ABC1.2` becomes `ABC`). Genes that cannot be matched through any of these steps are flagged for removal.

In addition to our custom CTD-based gene-matching algorithm, we applied gene normalization using NCBI Gene, MirGeneDB, HGNC, and UniProt. NCBI Gene was used to resolve gene identifiers, official symbols, names, and synonyms; MirGeneDB supported microRNA normalization; HGNC provided approved human gene symbols and aliases; and UniProt supported gene identification via protein names and aliases. Species information was determined using species-specific naming conventions. When species could not be determined, ‘human’ was used as the default since CTD also uses human identifiers when available.

First, we check for microRNA mentions; MirGeneDB is prioritized to identify precursor loci, which are then linked to NCBI Gene identifiers. Then, for other gene mentions, normalization first attempts NCBI symbol matching, followed by combined NCBI/HGNC synonym matching, UniProt protein-name matching, and our custom gene matching function. Gene names are used as a fallback when gene symbols cannot be resolved. Results of gene normalization can be found in Appendix S5.

**Interaction types:** Each extracted interaction is checked against a fixed list of 47 valid CTD interaction types and 3 valid prefixes (affects, increases, or decreases). Any interaction that does not match this list is either corrected to the closest valid type that is still supported by the source text or flagged for removal.

**Filtering:** Once all three standardization steps are complete, a final round of filtering is applied. Duplicate interactions are removed, with preference given to those extracted from structured sources such as tables and supplements over plain text and figures. We further remove any flagged entities and interactions from the standardization steps. Often, the LLM outputs include the same chemical-gene pair with the same interaction type but a different prefix, like `affects expression` and `increases expression`. Here, we keep the specific interaction like `increases expression` over `affects expression`.

Further, we remove interactions where the source or the data component indicates prior work or common knowledge (e.g., *Chemical X is known to downregulate gene Y*), negative interactions, and/or non-significance between the chemical and gene. Interactions from discussion sections are removed if the source text contains citation signals such as ‘‘et al.’’, ‘‘reported that’’, or ‘‘previous studies’’. Interactions are dropped if the source text contains speculative language (e.g., ‘‘might’’, ‘‘possibly’’, ‘‘previously shown’’) or broad generalizations (e.g., ‘‘it is known’’, ‘‘widely believed’’, ‘‘according to studies’’). Negative and null findings are identified and removed using a list of phrases such as ‘‘no effect’’, ‘‘did not alter’’, ‘‘not significant’’, and ‘‘failed to’’. Additionally, source text containing inline citation brackets (e.g., `[1]`) is removed as a further indicator of prior-work references. We use prior work identifying speculative and uncertain language in biomedical text to define our filtering patterns [1, 2]. Any remaining rows where the chemical, gene, or interaction could not be standardized are dropped. What remains is a clean, deduplicated set of chemical-gene interactions.

If the interaction stems from a discussion-type paragraph, we remove the interaction if the discussion contains citation signals indicative of prior work, including citations in brackets, ”et al.”, ”reported that”, ”according to”, ”previous studies”, ”previous works”, ”author”, or ”authors”.

Additionally, if the LLM source output contains speculative, uncertain, or prior-work language, including hedging terms (e.g., ”might”, ”possibly”, ”potentially”), generalization signals (e.g., ”known”, ”established”, ”documented”), and references to other or previous work (e.g., ”previously”, ”reported”, ”studied”).

Fig 2 displays a list of patterns we use to identify and remove interactions [1, 2] where the interaction is negative or null between the entities. Additionally, Fig 3 shows patterns of broad generalizations or common knowledge statements, including consensus language. If an interaction source text contains these broad patterns, it is removed.

Filtering relies on predefined textual patterns rather than a complete interpretation of experimental context. Thus, valid interactions may be excluded when their source text contains language associated with negative or null findings.

**Negative and Null Filtering Patterns**

**No effect / lack of effect:** lack of effect, no effect, without effect, did not affect, does not affect, did not alter, does not alter, did not induce, does not induce, did not change, does not change, was not affected, was not altered, was not induced, had no effect, no observable effect

**Negative results / failed outcomes:** failed to, unable to, cannot conclude, did not demonstrate, did not observe, was ineffective, no significant change, no significant effect, not significant, insignificant

**Lack of detection:** not detectable, undetectable, could not detect, cannot detect

**Contradictory associations:** not associated with, no correlation, no relationship, did not correlate, was not linked, not linked, no impact, did not impact, no influence, did not influence

**Unconfirmed findings:** did not support, failed to support, was not confirmed, cannot confirm, did not confirm, not reproducible, irreproducible

**Fig 2.** Negative and null filtering patterns used in the Refinement stage to remove interactions where the source text indicates no effect, failed outcomes, undetectable measurements, contradictory associations, or unconfirmed findings. These patterns do not automatically exclude decreases in measurable gene activity or expression.

**Broad and General Statement Filtering Patterns**

**Consensus / common knowledge:** generally, commonly, in general, common, general consensus, it is known, widely believed, well established, generally accepted, widely recognized, extensively studied

**Hedged associations:** appears to, tends to, is associated with, it is thought, it has been suggested, considered to be

**Prior literature references:** previous research, has been reported, according to studies, commonly observed, studies show, reported previously, as demonstrated, numerous studies

**Fig 3.** Broad and general statement filtering patterns used in the Refinement stage to remove interactions where the source text reflects common knowledge, hedged associations, or references to prior literature rather than direct experimental evidence.

### S5 Appendix. Prompt Design for CLOVER

Prompt engineering for this study involved several rounds of iterative refinement to develop the current prompt design. Given that the task is complex and the prompts are large, we structured them clearly for the LLM to understand the requirements. Here, we explain the key design choices we made and the LLM threats mitigated for the prompts.

#### Key Design Choices

**System Prompt:** The system prompt is the initial point of interaction with the LLM. Here, we state the goal, tone, and some conditions to limit the LLM's behavior [3]. We treat the LLM as a task-specific tool for extracting or validating chemical-gene interactions.

For the extraction prompt, we set clear instructions for the LLM to be truthful, emphasizing that if it is uncertain, it should admit its lack of knowledge rather than fabricate interactions. Additionally, we reiterate the goal and prohibit the LLM from introducing information not present in the passage.

**Structured Formatting:** We adopt a structured format [4] to present the LLM tasks and organize the prompts using headers, numbered lists, and bullet points. This structured approach allows the LLM to parse large prompts and efficiently reduce ambiguity. Visual cues (e.g., bold text) enhance clarity for important instructions. Each section of our prompt is demarcated to guide the LLM step by step, which mitigates misinterpretation risk. In addition, we provide an example of a structured return format, CSV, which ensures that responses are easily interpretable and usable in downstream processes.

**Clarity and Conciseness:** We define clear and direct instructions for our tasks, which are enhanced through structured formatting. We also implement strict rules to define what the LLM can and cannot do, and reiterate key points to maintain a straightforward process [5].

**One-Shot and Few-Shot learning:** We use one-shot learning with an example of a passage and extracted interactions. For validation, we use few-shot learning [6] with two examples of validated interactions. The provided examples cover various scenarios and demonstrate the type of output we expect. One-Shot and Few-Shot learning help guide the LLM toward the correct format and response without overwhelming the LLM with excessive data.

### LLM Threats and Mitigation Strategies

**''Lost in the Middle'' Phenomenon:** The ''Lost in the Middle'' phenomenon is when an LLM fails to encompass all aspects of a complex prompt in the response. To mitigate this possibility, we use structured formatting and place key information at the beginning or end of the prompt. For example, in our study, the Extraction LLM failed to identify the correct interaction from a list of 47 possible interactions. By placing the interaction list earlier in the prompt, the LLM's ability to identify the proper interaction improved.

**Spurious Associations:** Relation extraction using LLMs introduces the spurious associations threat. Spurious associations occur when the LLM incorrectly links entities based on an unobserved factor rather than a genuine causal interaction. Since LLMs are trained on vast amounts of text, they can perform poorly on unseen data or when underlying assumptions change. In our work, spurious associations can arise in the Extraction LLM stage. We mitigate this risk in two ways: (i) by using PubTator to ground chemical and gene mentions to standardized CTD identifiers rather than relying on the extraction LLM to identify entities, and (ii) by applying rule-based corrections in Refinement to eliminate interactions that do not conform to CTD classification standards, filtering out extractions that lack sufficient grounding in the source text.

**Hallucinations:** LLM Hallucinations occur when the LLM generates responses not supported by the input text, potentially leading to false and nonsensical outputs. We address hallucinations by enforcing strict rules and requiring the LLM to explicitly reference the specific sections of the passage text for each interaction. Furthermore, we instruct the LLM to state when it does not have sufficient information rather than hallucinating.

**LLM Deterministic Traits:** We set the temperature to zero to reduce randomness and set the seed to 42 to enforce the deterministic behavior of LLMs. We did not set top_p or top_k, as these parameters and temperature influence output randomness [7]. Setting all parameters can introduce unpredictable and potentially contradictory behavior.

**S6 Appendix.** Additional Tables and Figures

Fig 4 is a comprehensive multi-perspective count of extraction quality across all 18 model configurations (single-model and chained variants) at the Standardized and Filtered stages. We use exact matching between predicted and CTD-curated interactions based on PMID, chemical ID, gene symbol, and interaction prefix and type. A prediction is counted as a true positive only when all fields match. Predictions with the correct chemical-gene pair but an incorrect interaction are classified as partial matches for error analysis only and count as false positives. Partial matches receive no credit in precision or recall calculations.

Fig 4a is a stacked bar chart showing the count of correct, partial-match, and incorrect interactions extracted by each model after CTD-standardization and filtering steps. Fig 4b is also a stacked bar chart showing the same counts as Fig 4a but with no filtering. Solid bars represent true positives, hatched bars represent partial matches, and the light bars are incorrect in all aspects of the interaction. Table sections are the largest source of interactions but also the largest source of errors because they cannot be filtered due to a lack of context. Figure components are the smallest in volume but have high error rates due to the absence of PubTator NER for image content.

Fig 4c reports five classes of entity and interaction errors in the standardized and filtered outputs across all configurations. Chemicals and genes are marked as errors if absent from CTD for the 50 PMC articles. Interactions are marked as errors if they have the correct chemical-gene pair, but either the interaction type, prefix, or type and prefix are invalid.

**Invalid Entities** Invalid genes are the most prevalent error class and occur when the output has associated the gene with an incorrect chemical, interaction, or is not of interest to CTD. Invalid genes can also occur from the gene matching algorithm described in Materials and Methods. Invalid chemicals follow a similar pattern to invalid gene symbols but are less frequent because they are better identified by PubTator NER. Figure data is more susceptible to invalid entities because of their unstructured nature.

**Table 1.** Article-level extraction performance and processed document components for the 50 evaluated articles. Sec. is the number of data components; CTD represents CTD-curated interactions; Ext. represents extracted interactions; Match represents matched interactions; P and R are precision and recall, respectively. Txt, Tbl, Fig, and Sup are processed text, table, figure, and supplement data components, respectively.

| PMID | PMCID | Sec. | CTD | Ext. | Match | P | R | Txt | Tbl | Fig | Sup |
|---|---|---|---|---|---|---|---|---|---|---|---|
| 22110480 | PMC3205718 | 22 | 2 | 2 | 2 | 1.000 | 1.000 | 16 | 1 | 5 | 0 |
| 34908893 | PMC8640676 | 19 | 1 | 1 | 1 | 1.000 | 1.000 | 11 | 3 | 5 | 0 |
| 29380240 | PMC6060758 | 27 | 7 | 16 | 7 | 0.438 | 1.000 | 20 | 3 | 4 | 0 |
| 15953866 | PMC2782200 | 19 | 1 | 3 | 1 | 0.333 | 1.000 | 13 | 0 | 6 | 0 |
| 32298299 | PMC7161975 | 101 | 1 | 3 | 1 | 0.333 | 1.000 | 78 | 0 | 23 | 0 |
| 17183729 | PMC1762369 | 76 | 5 | 32 | 5 | 0.156 | 1.000 | 8 | 1 | 4 | 63 |
| 18280460 | PMC2279149 | 39 | 3 | 22 | 3 | 0.136 | 1.000 | 26 | 4 | 9 | 0 |
| 22747577 | PMC3489685 | 28 | 158 | 160 | 154 | 0.962 | 0.975 | 13 | 2 | 8 | 5 |
| 24413757 | PMC3907846 | 27 | 35 | 48 | 34 | 0.708 | 0.971 | 17 | 2 | 8 | 0 |
| 22808131 | PMC3392256 | 38 | 90 | 87 | 82 | 0.943 | 0.911 | 23 | 8 | 7 | 0 |
| 15811834 | PMC1278483 | 38 | 8 | 10 | 7 | 0.700 | 0.875 | 28 | 0 | 10 | 0 |
| 22228119 | PMC3349910 | 23 | 112 | 113 | 97 | 0.858 | 0.866 | 17 | 2 | 4 | 0 |
| 17524151 | PMC1890552 | 29 | 5 | 14 | 4 | 0.286 | 0.800 | 22 | 0 | 7 | 0 |
| 18299717 | PMC2243244 | 15 | 76 | 104 | 59 | 0.567 | 0.776 | 8 | 6 | 1 | 0 |
| 27153807 | PMC4859981 | 29 | 4 | 3 | 3 | 1.000 | 0.750 | 20 | 0 | 9 | 0 |
| 16393664 | PMC1332662 | 10 | 61 | 46 | 41 | 0.891 | 0.672 | 2 | 3 | 5 | 0 |
| 33000210 | PMC7533447 | 28 | 6 | 12 | 4 | 0.333 | 0.667 | 23 | 1 | 4 | 0 |
| 19889203 | PMC3003513 | 68 | 6 | 55 | 4 | 0.073 | 0.667 | 56 | 0 | 12 | 0 |
| 25859307 | PMC4387225 | 35 | 8 | 9 | 5 | 0.556 | 0.625 | 17 | 3 | 15 | 0 |
| 27397436 | PMC5298042 | 72 | 222 | 178 | 134 | 0.753 | 0.604 | 19 | 1 | 4 | 48 |
| 23568217 | PMC3620667 | 26 | 7 | 5 | 4 | 0.800 | 0.571 | 21 | 0 | 5 | 0 |
| 30104959 | PMC6077207 | 34 | 21 | 18 | 10 | 0.556 | 0.476 | 26 | 0 | 8 | 0 |
| 21294877 | PMC3042928 | 29 | 8 | 13 | 3 | 0.231 | 0.375 | 23 | 0 | 6 | 0 |
| 19617202 | PMC3136680 | 27 | 3 | 5 | 1 | 0.200 | 0.333 | 19 | 0 | 8 | 0 |
| 25100957 | PMC4104784 | 22 | 49 | 17 | 14 | 0.824 | 0.286 | 14 | 3 | 5 | 0 |
| 17118201 | PMC1676006 | 28 | 4 | 3 | 1 | 0.333 | 0.250 | 23 | 0 | 5 | 0 |
| 24403256 | PMC3892387 | 37 | 4 | 17 | 1 | 0.059 | 0.250 | 32 | 0 | 5 | 0 |
| 22571318 | PMC3485091 | 33 | 6 | 10 | 1 | 0.100 | 0.167 | 26 | 0 | 7 | 0 |
| 20709644 | PMC2974693 | 35 | 13 | 17 | 1 | 0.059 | 0.077 | 31 | 3 | 1 | 0 |
| 19114014 | PMC2631511 | 16 | 1 | 1 | 0 | 0.000 | 0.000 | 7 | 0 | 9 | 0 |
| 20801540 | PMC2987498 | 26 | 1 | 6 | 0 | 0.000 | 0.000 | 18 | 1 | 7 | 0 |
| 22562770 | PMC3440069 | 27 | 3 | 6 | 0 | 0.000 | 0.000 | 24 | 0 | 3 | 0 |
| 23349674 | PMC3551951 | 32 | 1 | 2 | 0 | 0.000 | 0.000 | 25 | 2 | 5 | 0 |
| 27517156 | PMC5295416 | 16 | 1 | 1 | 0 | 0.000 | 0.000 | 10 | 0 | 6 | 0 |
| 27736846 | PMC5129817 | 28 | 1 | 3 | 0 | 0.000 | 0.000 | 22 | 4 | 2 | 0 |
| 28637469 | PMC5480112 | 11 | 1 | 3 | 0 | 0.000 | 0.000 | 7 | 4 | 0 | 0 |
| 30114287 | PMC6095593 | 7 | 10 | 1 | 0 | 0.000 | 0.000 | 2 | 1 | 4 | 0 |
| 26714306 | PMC4695093 | 10 | 1 | 0 | 0 | – | 0.000 | 6 | 3 | 1 | 0 |
| 31064381 | PMC6503545 | 16 | 2 | 0 | 0 | – | 0.000 | 11 | 0 | 5 | 0 |
| 37047793 | PMC10095583 | 6 | 4 | 0 | 0 | – | 0.000 | 2 | 1 | 3 | 0 |
| 19654922 | PMC2717139 | 22 | 0 | 2 | 0 | 0.000 | – | 13 | 2 | 7 | 0 |
| 22112310 | PMC3279445 | 21 | 0 | 3 | 0 | 0.000 | – | 15 | 4 | 2 | 0 |
| 23006841 | PMC3488501 | 31 | 0 | 3 | 0 | 0.000 | – | 26 | 0 | 5 | 0 |
| 24039778 | PMC3769399 | 50 | 0 | 8 | 0 | 0.000 | – | 32 | 2 | 16 | 0 |
| 26039251 | PMC4454644 | 6 | 0 | 6 | 0 | 0.000 | – | 2 | 1 | 3 | 0 |
| 26492236 | PMC4632778 | 33 | 0 | 9 | 0 | 0.000 | – | 20 | 1 | 12 | 0 |
| 31378678 | PMC6706627 | 104 | 0 | 26 | 0 | 0.000 | – | 16 | 1 | 7 | 80 |
| 36638633 | PMC10016736 | 45 | 0 | 15 | 0 | 0.000 | – | 43 | 2 | 0 | 0 |
| 25361045 | PMC4245613 | 16 | 0 | 0 | 0 | – | – | 10 | 1 | 5 | 0 |
| 29045497 | PMC5646803 | 26 | 0 | 0 | 0 | – | – | 18 | 1 | 7 | 0 |

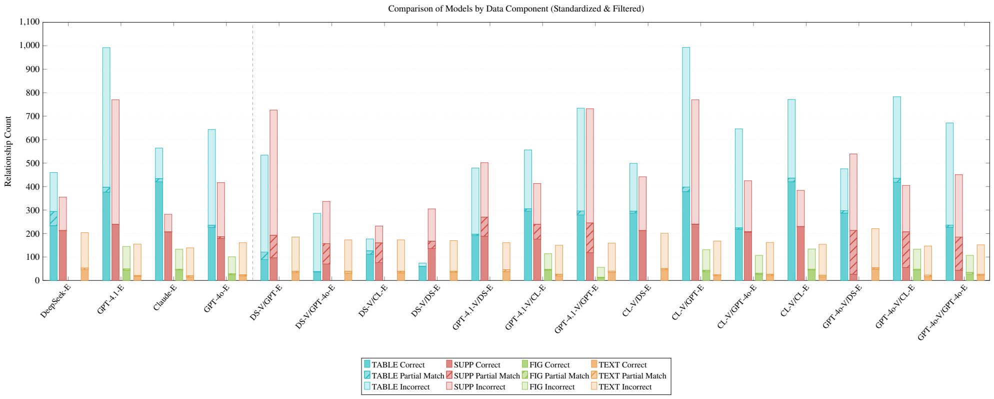


**Fig. 8a. Standardized & Filtered stacked bar chart.** Standardized & Filtered stacked bar chart showing the count of correct (solid), partial match (hatched), and incorrect (transparent) interactions per data component (TABLE, SUPPLEMENT, FIGURE, TEXT) after CTD-standardization and filtering steps.

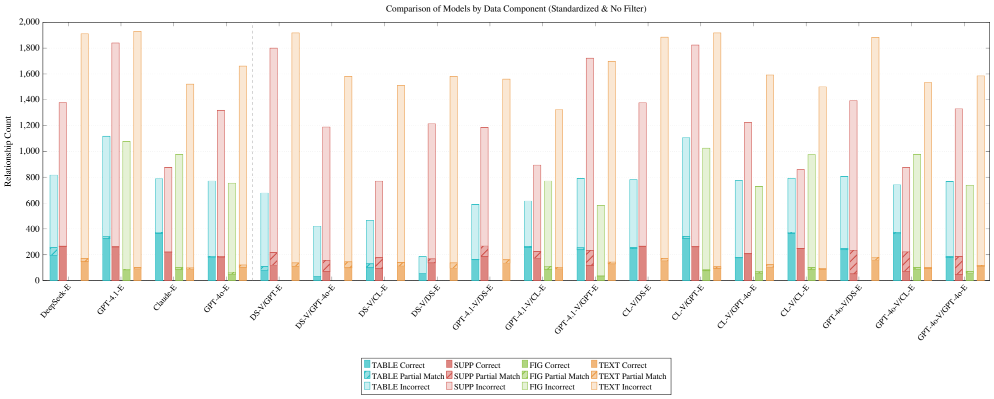


**Fig. 8b. Standardized-only stacked bar chart.** Standardized-only stacked bar chart (duplicates removed with structured-data priority, no filtering) showing the substantially larger candidate set produced before filtering, illustrating how filtering reduces the candidate interaction set.

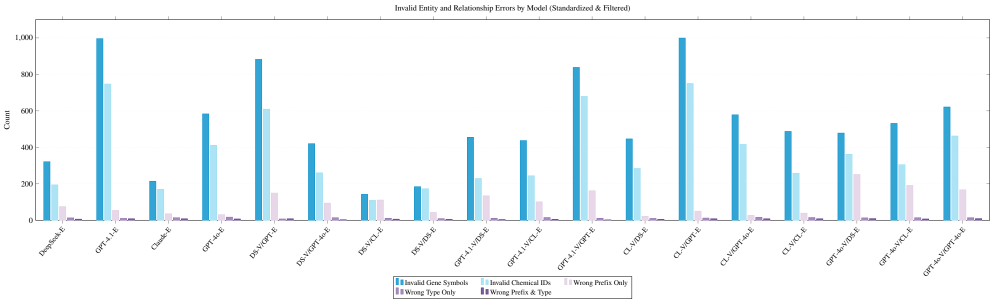


**Fig. 8c. Invalid entity and interaction error counts.** Invalid entity and interaction error counts per configuration at the Standardized & Filtered stage, decomposed into five error classes: invalid gene symbols, invalid chemical IDs, wrong prefix only, wrong interaction type only, and wrong prefix and type combined. Invalid gene symbols are the most prevalent error class, reflecting the difficulty of resolving gene identifiers from informal or ambiguous mentions in biomedical text. Error categories overlap; therefore, the sum of errors cannot be used to obtain the total number of incorrect predictions. CL-E produces fewer prefix-only errors than other models. Validation does not reduce interaction errors, supporting the finding that LLM chaining is not an effective strategy for this task.

**Fig 4. Relationship count.** Relationship counts for filtered and CTD-standardized (Fig. 4a) and CTD-standardized (Fig. 4b) and entity/interaction error analysis (Fig. 4c) across all 18 model configurations.

**Table 2.** Changes in CTD-matched (TP) and CTD-unmatched (FP) interactions following same-model validation for standardized and filtered output. Retained interactions occur in both stages; lost and removed interactions disappear after validation; gained and introduced interactions appear after validation. Changes in precision ($\Delta P$) and recall ($\Delta R$) are reported in percentage points relative to standalone extraction.

| Model | TP Ret. | TP Rem. | TP Gain | FP Ret. | FP Rem. | FP Gain | $\Delta P$ (%) | $\Delta R$ (%) |
|---|---|---|---|---|---|---|---|---|
| DeepSeek | 175 | 316 | 54 | 285 | 243 | 35 | -6.47 | -27.52 |
| GPT-4.1 | 440 | 235 | 2 | 1069 | 318 | 170 | -6.44 | -24.47 |
| Claude | 684 | 0 | 24 | 427 | 7 | 308 | -12.12 | +2.52 |
| GPT-4o | 313 | 135 | 4 | 738 | 136 | 326 | -10.93 | -13.76 |

**Invalid interaction type, prefix, or prefix & type** Wrong interaction prefix errors are informative because the LLM successfully identifies the correct chemical, gene, and interaction type but fails to determine directionality. CL-E produces fewer prefix errors, which is consistent with its tendency towards more accurate directional assignment and higher instruction fidelity. Wrong type and prefix & type errors occur less frequently than wrong prefix errors. These occur typically when the LLM cannot decide between activity and expression and chooses the incorrect interaction, especially in figure interactions. Validation does not reduce these errors, further supporting that chaining is not an effective method for this task.

Fig 5 shows the stacked distributions of interaction types of all model configurations compared to the CTD-curated reference interactions for the 50 PMC articles. CTD distribution is concentrated on *increases* and *decreases expression*, which is also reflected in the model extractions. Even though less common interaction types are represented in the LLM output, the models tend to choose to *increase* or *decrease expression* because the rare interactions often appear in highly specific experimental contexts that the LLM may not associate with a formal CTD interaction type. Conversely, a curator and a model may decide on different rare interactions, yet both assignments can be justifiable depending on the interpretative perspective and the specific biological context considered. For example, authors in a paper may say a particular chemical increases the activation of a gene, and further in the paper, they may describe another relationship. The LLM would extract both because it views the relationship from the specific context it has. A curator, depending on the context, may select one or both of the relationships described in the context.

In our evaluation set of 50 articles, 17 interactions were represented in the corresponding CTD data, reflecting the inherent nature of the chemical–gene interaction data in CTD. Expression-related interactions account for 95% of the 952 chemical-gene interactions curated by CTD. Table 3 provides the class-level distribution of CTD-curated interactions and Claude predictions. Claude succeeded in capturing seven unique interaction classes, most of which are related to gene expression. However, Claude extracted 29 different interaction classes, and out of these, 17 classes were not included in CTD. The relationship imbalance in our evaluation set is also reflected in the broader distribution of CTD-curated interactions, where expression-related interactions account for 85% of all CTD single-step relationships.

Table 4 summarizes gene normalizations made using the custom CTD-based gene matching function and four external reference databases. We manually analyzed potential gene normalization errors for CL-E outputs. We analyzed a random set of 100 gene normalizations out of 624 normalized CL-E rows. Potential incorrect mappings included MEK to MAP2K7, SOD to SOD1, SP to TFF2, and MIR3545 to MIR203.

For CL-E, filtering reduced the number of predictions from 4,121 to 1,118 (Table 5, increasing CTD-agreement precision from 18% to 61%, and recall decreased from 79% to 72%. Of the 729 unique CTD-matched relationships present before filtering, 45 were lost. Most lost relationships involved changes in gene expression. These findings indicate that rule-based filtering improves precision but can also exclude CTD-supported interactions.

**S7 Appendix.** Implementation Details

The Clover pipeline processes PubMed Central full-text articles through a fully automated process (excluding the final Curator Review step, which is human review). Everything, including the webscrapers, is implemented in Python. The LLMs are employed via API calls. On average, the pipeline takes 7-10 minutes per article. The logical flow is as follows:

- **Article Download and Preprocessing** (1 minute per article). This step consists of retrieving a full-text article from PubMed Central using PubMed ID, downloading the article from PubTator, using the webscraper to extract

**Table 3.** The table compares CTD-curated interactions with CL-E predictions after standardization and filtering. TP represents exact CTD-matched predictions; Unmatched represents predictions without an exact CTD match. P and R denote precision and recall against CTD.

| **Relationship** | **CTD** | **Claude** | **TP** | **Unmatched** | **P (%)** | **R (%)** |
|---|---|---|---|---|---|---|
| increases expression | 540 | 528 | 397 | 131 | 75.2 | 73.5 |
| decreases expression | 351 | 362 | 265 | 97 | 73.2 | 75.5 |
| decreases activity | 12 | 60 | 8 | 52 | 13.3 | 66.7 |
| affects expression | 12 | 32 | 0 | 32 | 0.0 | 0.0 |
| increases secretion | 10 | 11 | 7 | 4 | 63.6 | 70.0 |
| increases activity | 8 | 36 | 3 | 33 | 8.3 | 37.5 |
| affects response to substance | 4 | 0 | 0 | 0 | – | 0.0 |
| decreases response to substance | 4 | 0 | 0 | 0 | – | 0.0 |
| increases cleavage | 3 | 3 | 3 | 0 | 100.0 | 100.0 |
| decreases secretion | 1 | 4 | 0 | 4 | 0.0 | 0.0 |
| increases acetylation | 1 | 4 | 0 | 4 | 0.0 | 0.0 |
| affects localization | 1 | 2 | 0 | 2 | 0.0 | 0.0 |
| increases localization | 1 | 2 | 1 | 1 | 50.0 | 100.0 |
| increases methylation | 1 | 1 | 0 | 1 | 0.0 | 0.0 |
| affects abundance | 1 | 0 | 0 | 0 | – | 0.0 |
| decreases export | 1 | 0 | 0 | 0 | – | 0.0 |
| increases response to substance | 1 | 0 | 0 | 0 | – | 0.0 |
| affects binding | 0 | 18 | 0 | 18 | 0.0 | – |
| affects cotreatment | 0 | 10 | 0 | 10 | 0.0 | – |
| decreases phosphorylation | 0 | 8 | 0 | 8 | 0.0 | – |
| affects activity | 0 | 7 | 0 | 7 | 0.0 | – |
| decreases uptake | 0 | 6 | 0 | 6 | 0.0 | – |
| decreases cleavage | 0 | 4 | 0 | 4 | 0.0 | – |
| decreases localization | 0 | 4 | 0 | 4 | 0.0 | – |
| affects methylation | 0 | 3 | 0 | 3 | 0.0 | – |
| decreases degradation | 0 | 3 | 0 | 3 | 0.0 | – |
| decreases oxidation | 0 | 2 | 0 | 2 | 0.0 | – |
| increases stability | 0 | 2 | 0 | 2 | 0.0 | – |
| affects metabolic processing | 0 | 1 | 0 | 1 | 0.0 | – |
| affects splicing | 0 | 1 | 0 | 1 | 0.0 | – |
| decreases hydroxylation | 0 | 1 | 0 | 1 | 0.0 | – |
| decreases metabolic processing | 0 | 1 | 0 | 1 | 0.0 | – |
| decreases ubiquitination | 0 | 1 | 0 | 1 | 0.0 | – |
| increases uptake | 0 | 1 | 0 | 1 | 0.0 | – |
| **Total** | **952** | **1,118** | **684** | **434** | **61.2** | **71.8** |

**Table 4.** Gene normalization across the 18 CLOVER model configurations. Rows is the number of LLM predictions for each standardized and filtered configuration. The remaining columns report gene-field modifications attributed to the custom CTD-based matching function and normalization using MirGeneDB, NCBI Gene, HGNC, and UniProt. Counts indicate modifications, not verified corrections.

| Model | Rows | Custom CTD | MirGeneDB | NCBI Gene | HGNC | UniProt |
|---|---|---|---|---|---|---|
| DS-E | 1,019 | 119 | 85 | 268 | 24 | 0 |
| DS-V/DS-E | 549 | 11 | 133 | 152 | 13 | 0 |
| GPT-4.1-V/DS-E | 1,142 | 58 | 139 | 378 | 25 | 1 |
| CL-V/DS-E | 1,142 | 121 | 85 | 376 | 24 | 0 |
| GPT-4o-V/DS-E | 1,236 | 125 | 109 | 408 | 25 | 1 |
| GPT-4.1-E | 2,062 | 135 | 130 | 891 | 42 | 2 |
| DS-V/GPT-4.1-E | 1,445 | 82 | 132 | 660 | 9 | 2 |
| GPT-4.1-V/GPT-4.1-E | 1,681 | 82 | 119 | 742 | 31 | 0 |
| CL-V/GPT-4.1-E | 2,062 | 135 | 130 | 887 | 41 | 2 |
| GPT-4o-E | 1,322 | 61 | 65 | 518 | 7 | 0 |
| DS-V/GPT-4o-E | 796 | 21 | 56 | 308 | 1 | 0 |
| CL-V/GPT-4o-E | 1,340 | 71 | 65 | 522 | 8 | 0 |
| GPT-4o-V/GPT-4o-E | 1,381 | 71 | 64 | 550 | 8 | 0 |
| CL-E | 1,118 | 72 | 120 | 422 | 20 | 2 |
| DS-V/CL-E | 582 | 27 | 120 | 136 | 8 | 2 |
| GPT-4.1-V/CL-E | 1,233 | 52 | 128 | 532 | 13 | 0 |
| CL-V/CL-E | 1,443 | 81 | 117 | 601 | 20 | 2 |
| GPT-4o-V/CL-E | 1,468 | 84 | 121 | 617 | 18 | 2 |

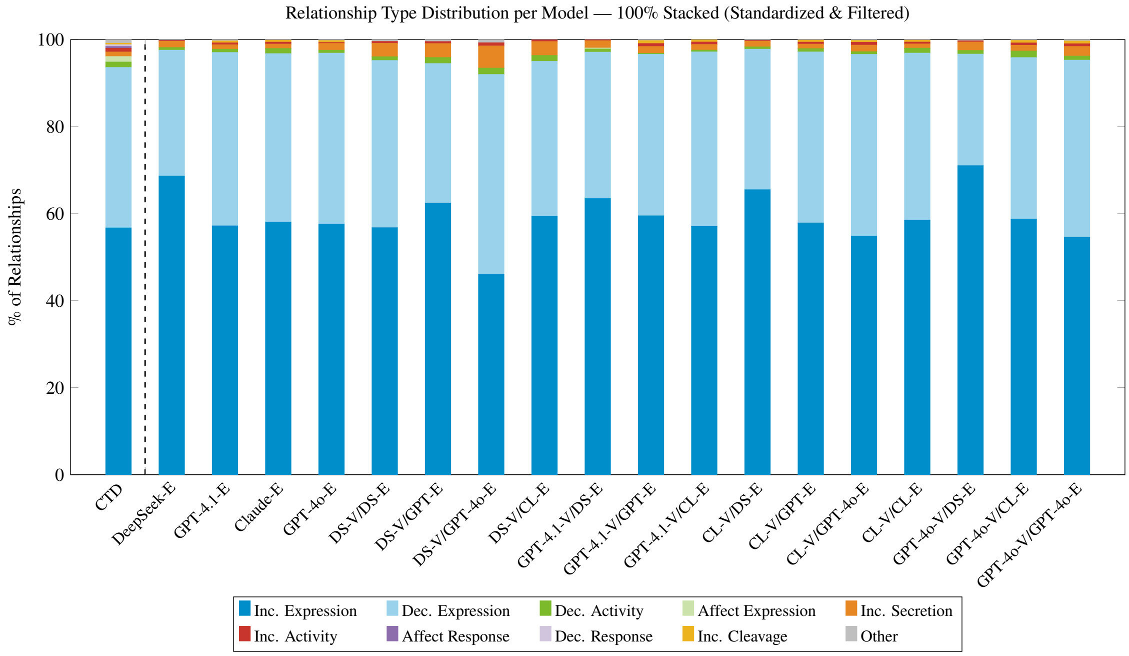


**Fig 5. Relationship Type Distribution per Model.** Stacked bar chart comparing the distribution of interaction types across all 18 CLOVER model configurations and CTD-curated relationships for the 50 PMC articles, shown at the Standardized & Filtered stage. Relationship types are color-coded, with the most frequent CTD-curated types highlighted. CTD-curated relationship distribution is heavily concentrated in expression-related interactions, a pattern broadly reflected across model outputs. Model distributions match CTD exact-matched chemical, gene, and interaction relationships.

**Table 5.** Effect of rule-based filtering across CLOVER configurations. Pre-filter and post-filter denote the number of standardized predictions before and after filtering. CTD-matched lost represents CTD-matched predictions removed during filtering; CTD-unmatched removed represents filtered predictions without a CTD match.

| Model | Pre-filter | Post-filter | Removed | CTD-matched lost | CTD-unmatched removed |
|---|---|---|---|---|---|
| *Standalone Extraction* | | | | | |
| DS-E | 4,049 | 1,019 | 3,030 | 58 | 2,972 |
| GPT-4.1-E | 5,938 | 2,062 | 3,876 | 51 | 3,825 |
| CL-E | 4,121 | 1,118 | 3,003 | 45 | 2,958 |
| GPT-4o-E | 4,497 | 1,322 | 3,175 | 56 | 3,119 |
| *DeepSeek Extractor* | | | | | |
| DS-V/DS-E | 2,974 | 549 | 2,425 | 55 | 2,370 |
| GPT-4.1-V/DS-E | 3,328 | 1,142 | 2,186 | 58 | 2,128 |
| CL-V/DS-E | 3,986 | 1,142 | 2,844 | 61 | 2,783 |
| GPT-4o-V/DS-E | 4,055 | 1,236 | 2,819 | 60 | 2,759 |
| *GPT-4.1 Extractor* | | | | | |
| DS-V/GPT-4.1-E | 4,369 | 1,445 | 2,924 | 66 | 2,858 |
| GPT-4.1-V/GPT-4.1-E | 4,793 | 1,681 | 3,112 | 70 | 3,042 |
| CL-V/GPT-4.1-E | 5,845 | 2,062 | 3,783 | 52 | 3,731 |
| *Claude Extractor* | | | | | |
| DS-V/CL-E | 2,722 | 582 | 2,140 | 60 | 2,080 |
| GPT-4.1-V/CL-E | 3,600 | 1,233 | 2,367 | 63 | 2,304 |
| CL-V/CL-E | 4,084 | 1,443 | 2,641 | 45 | 2,596 |
| GPT-4o-V/CL-E | 4,085 | 1,468 | 2,617 | 50 | 2,567 |
| *GPT-4o Extractor* | | | | | |
| DS-V/GPT-4o-E | 3,188 | 796 | 2,392 | 62 | 2,330 |
| CL-V/GPT-4o-E | 4,310 | 1,340 | 2,970 | 63 | 2,907 |
| GPT-4o-V/GPT-4o-E | 4,414 | 1,381 | 3,033 | 68 | 2,965 |

supplements and images, splitting the article into sections: main text (paragraphs), structured tables, figures, and supplements along with captions if provided, and creating a CSV file that has the sections with the appropriate prompts.

- **LLM Extraction** (2 to 3 minutes per article). Submit prompts to the target LLM API (GPT-4.1, GPT-4o, Claude-Sonnet-4.5, or DeepSeek-V4.1-Flash) with temperature=0.0. Parse CSV-formatted responses and aggregate per article. DeepSeek-V4.1-Flash takes the longest, around 12-15 minutes per article.
- **LLM Validation** (2 to 3 minutes per article). Construct the validation prompts. Then, submit the prompts to a second LLM using the target LLM API with temperature=0.0. DeepSeek-V4.1-Flash takes the longest, around 12-15 minutes per article.
- **Rule-Based Refinement** (1 to 2 minutes per article). Extract the LLM output and process it into a CSV table. Most of the time here is taken to read CTD data files and construct lookup tables for chemical and gene entities. Once the data is read and lookup tables are constructed, standardization and filtering take around 20 seconds per article. After this, the output is a clean CSV table ready for curator review.

Now, we consider the hardware computational infrastructure and technical environment.

- **LLM API calls:** All extraction and validation stages rely on commercial LLM APIs (OpenAI GPT-4.1/GPT-4o, Anthropic Claude, DeepSeek-V4.1-Flash). No on-premises GPU or local model hosting was used.
- **Data pre- and postprocessing:** Preprocessing and rule-based refinement are implemented as lightweight Python scripts (CPU-bound, no GPU required), executable on standard compute servers or laptops.
- **Storage:** Extracted CSV outputs (10-50 KB per article after filtering) are stored locally.

Cost considerations for a 50-article benchmark are detailed below. These models also offer token caching (saving tokens when the same tokens are repeated) and batch processing for higher-level subscriptions, which can reduce token usage because most prompts contain repeated information.

We show the cost per prompt in Table 6. In this work, we tested using 1,563 prompts. The input cost range is represented using a 1,500-70,000 token range per prompt with no token caching and no batch processing. The output cost is an average of the 50 articles processed in this work, which is 200 tokens.

| Model | Input Cost ($) | Output Cost ($) |
|---|---|---|
| GPT-4.1 | 0.003 – 0.14 | 0.0016 |
| GPT-4o | 0.004 – 0.175 | 0.002 |
| Claude-Sonnet-4.5 | 0.0045 – 0.21 | 0.003 |
| DeepSeek-V4.1-Flash | 0.00042 – 0.0196 | 0.000084 |

**Table 6.** Cost comparison of models (input and output pricing) per prompt.

Every month, CTD processes around 350 articles. Since we tested 50 articles with 1,563 sections, each article has an average of 31.26 sections. Then, 350 articles means 10.941 sections, bringing the total input tokens to 197 million. The input cost for Claude (most expensive) ranges from $49.23-$2,297.61 and the output cost is $32.82. Clover is currently implemented as standalone Python code and is not yet integrated into CTD's curation pipeline.

Experiments were run between October 12th and October 16th of 2026. The model identifiers are as follows: Claude-Sonnet-4.5 (claude-sonnet-4-5-20250929); GPT-4.1 (gpt-4.1-2025-04-14), GPT-4o (gpt-4o-2024-11-20); DeepSeek-V4.1-Flash (DeepSeek-V4.1-Flash-1017).

**S8 Fig.** Baseline Prompt (Fig 6).

**Naive Baseline Prompt using Zero-Shot Learning**

**System Prompt:**
You are an AI assistant that extracts relationships between chemicals and genes from scientific text.

**User Prompt:**
Extract all relationships between chemicals and genes from the text below. Format the output as CSV with one relationship per row.

**Columns:** `explanation`, `chemicalName`, `relationship`, `geneName`, `source`

The relationship must follow this format: `increase <type>`, `decrease <type>`, or `affect <type>`
Choose from the following relationship types:
List of relationships from Fig 5

Output only the CSV with no additional commentary.

**Fig 6. Naive baseline prompt.** Naive baseline prompt applied directly to full-text PDFs. Unlike CLOVER, this prompt provides no PubTator NER, no section splitting, no entity grounding, and no examples.

**S9 Appendix.** Extraction Prompts: Fig 7, Fig 8, Fig 9, Fig 10, Fig 11, Fig 12, Fig 13, and Fig 14.

**Fig 7.** System prompt for figure extraction.

**Fig 8.** User prompt for figure extraction.

**Fig 9.** System prompt for table extraction.

**Fig 10.** User prompt for table extraction. LLMs perform better with raw XML tables because tables are usually complex (may include multiple headers), and these do not parse properly.

**Fig 11.** System prompt for supplement extraction.

**Fig 12.** User prompt for supplement extraction.

**Fig 13.** System prompt for text extraction.

**Fig 14.** User prompt for text extraction.

**S10 Appendix.** Validation Prompts: Fig 15, Fig 16, Fig 17, Fig 18, Fig 19, Fig 20, Fig 21, and Fig 22.

**Fig 15.** System prompt for figure validation.

**Fig 16.** Structured validation prompt for figure data using one-shot learning.

**Fig 17.** System prompt for table validation.

**Fig 18.** Structured validation prompt for table data using one-shot learning.

**Fig 19.** System prompt for supplement validation.

**Fig 20.** Structured validation prompt for CSV supplement data using one-shot learning.

**Fig 21.** System prompt for text validation.

**Fig 22.** Structured validation prompt for text data using one-shot learning.

### References


1. Farkas R, Vincze V, Móra G, Csirik J, Szarvas G. The CoNLL-2010 Shared Task: Learning to Detect Hedges and Their Scope in Natural Language Text. In: Farkas R, Vincze V, Szarvas G, Móra G, Csirik J, editors. Proceedings of the Fourteenth Conference on Computational Natural Language Learning – Shared Task. Uppsala, Sweden: Association for Computational Linguistics; 2010. p. 1-12. Available from: `https://aclanthology.org/W10-3001/`.
2. Vincze V, Szarvas G, Farkas R, Móra G, Csirik J. The BioScope Corpus: Biomedical Texts Annotated for Uncertainty, Negation and Their Scopes. BMC Bioinformatics. 2008;9(Suppl 11):S9. doi:10.1186/1471-2105-9-S11-S9.
3. Giray L. Prompt Engineering with ChatGPT: A Guide for Academic Writers. Annals of Biomedical Engineering. 2023;51(12):2629-33.
4. He J, Rungta M, Koleczek D, Sekhon A, Wang FX, Hasan S. Does Prompt Formatting Have Any Impact on LLM Performance? arXiv preprint arXiv:241110541. 2024.
5. Sahoo P, Singh AK, Saha S, Jain V, Mondal S, Chadha A. A Systematic Survey of Prompt Engineering in Large Language Models: Techniques and Applications. arXiv preprint arXiv:240207927. 2024.
6. Brown T, Mann B, Ryder N, Subbiah M, Kaplan JD, Dhariwal P, et al. Language Models Are Few-Shot Learners. Advances in Neural Information Processing Systems. 2020;33:1877-901.
7. Peeperkorn M, Kouwenhoven T, Brown D, Jordanous A. Is Temperature the Creativity Parameter of Large Language Models? arXiv preprint arXiv:240500492. 2024.
