## Supplementary material for "CLOVER: Chemical-gene LLM benchmark for obtaining valid entity relationships for the Comparative Toxicogenomics Database (CTD)": User prompt for figure extraction.

**Structured Extraction Prompt for Figures using One-Shot Learning**

### Extract Specific Relationships Between Chemicals and Genes from image

#### Image Caption:
Cytotoxicity of SM on SAE and BTE cells. Cells were exposed to various concentrations of SM. Cell viability was measured using the MTS assay 24 h after exposure, and is expressed as a percentage of the control (unexposed cells, “C”). *** P < 0.001 versus control.

—

### Provided Relationship List
The following relationships are predefined and must be used to describe chemical-gene interactions:
List of relationships from Fig 5

—

### Instructions
For each extracted relationship, generate a CSV object with the following fields:

- `chemicalName`: Name of the chemical.
- `relationship`: This must be chosen from the provided list of relationships. The relationship may be prefixed by `increase`, `decrease`, or `affect`.
- `geneName`: Gene name if provided, otherwise leave blank. Gene names are longer than gene symbol.
- `geneSymbol`: Gene symbol if provided, otherwise leave blank. Gene symbols are typically all in uppercase.
- Do not infer relationships not supported by the provided content

#### Guidelines:
1. Extract specific relationships between these chemicals and genes using only the predefined terms from the provided relationship list. Do not use any terms or descriptions of interactions from the source text unless they exactly match terms from the list.
2. Provide only the CSV output—no additional explanations or comments.
3. Ensure that your CSV output is well-formed and follows CSV format.
4. Recheck to ensure the relationship is in the provided relationship list.
5. Use your best judgment to extract chemical-gene relationships; in some cases, there might not be any. If there are no relationships, leave the CSV empty.

—

### Example Output

#### Example Image Caption:
Gene regulation by Genistein.

#### Example Image:
Assume the image is passed in and you parsed the image internally.

#### Example CSV Output:

```
chemicalName,relationship,geneName,geneSymbol
"Genistein","Decreases expression","-","DEF2"
"Genistein","Increases expression","-","ABC1"
"Genistein","Decreases activity","tyrosine 3-monooxygenase","-"
```
