## Supplementary material for "CLOVER: Chemical-gene LLM benchmark for obtaining valid entity relationships for the Comparative Toxicogenomics Database (CTD)": User prompt for table extraction

**Structured Extraction Prompt for Tables using One-Shot Learning**

### Extract Specific Relationships Between Chemicals and Genes from the following table in XML format

#### Table Caption:
Genes up-regulated at least 1.5-fold in bisphenol A treatment in rats

#### Full Table:
```
<?xml version="1.0" encoding="UTF-8"?> <table frame="hsides" rules="groups">…[table content]…
```

#### Chemicals and their MESH IDs:
bisphenol A, C006780

#### Genes and their NCBI IDs:
- glucose-6-phosphate dehydrogenase, 855480
- glutathione peroxidase 3, 64317
- cytochrome b-5, 64001
- hydroxysteroid (17-beta) dehydrogenase 4, 79244
- collagen, type 1, alpha 1, 29393
- prostaglandin-endoperoxide synthase 1, 24693
- carbonyl reductase 1, 29224
- collagen, type III, alpha 1, 84032
- glutathione S-transferase, 58962

—

### Provided Relationship List
The following relationships are predefined and must be used to describe chemical-gene interactions: List of relationships from Fig 5

—

### Instructions
For each extracted relationship, generate a CSV object with the following fields:
- `chemicalName`: Name of the chemical
- `chemicalID`: Chemical ID if provided, otherwise leave blank.
- `relationship`: Must be chosen from the provided relationship list (may be prefixed by `increase`, `decrease`, or `affect`)
- `geneName`: Name of the gene
- `geneID`: Gene ID if provided, otherwise leave blank.

#### Guidelines:
1. Extract specific relationships included in the table.
2. Include ALL chemicals and genes in your output regardless of them being in the provided gene or chemical data. Set the ID field to `-` if not provided.
3. Provide only the CSV output—no additional explanations or comments.
4. Ensure that your CSV output is well-formed.
5. Recheck that the relationship is in the provided list.
6. In tabular data, relationships of interest are typically `increase expression` or `decrease expression` unless otherwise specified.
7. Do not infer relationships not supported by the provided content

—

### Example Output

```
chemicalName,chemicalID,relationship,geneName,geneID
"bisphenol A","C006780","increase expression","carbonyl
reductase 1","29224"
"bisphenol A","C006780","increase expression","glucose-6-phosphate dehydrogenase","855480"
```
