## Supplementary material for "CLOVER: Chemical-gene LLM benchmark for obtaining valid entity relationships for the Comparative Toxicogenomics Database (CTD)": User prompt for supplement extraction

**Structured Extraction Prompt for CSV Supplements using Few-Shot Learning**

### Extract Specific Relationships Between Chemicals and Genes from the following table in csv format with a header.

#### Table Caption:
Genes with a time-dependent, continuous increase or decrease in mRNA expression level during bisphenol A treatment.

#### Table Sheet Name:
ctr up >1.5

#### Full Table:
CSV Supplement content

—
### Provided Relationship List
The following relationships are predefined and must be used to describe chemical-gene interactions:
List of relationships from Fig 5

—
### Instructions
For each extracted relationship, generate a CSV object with the following fields:

- `chemicalName`: Name of the chemical.
- `relationship`: This must be chosen from the provided list of relationships. The relationship may be prefixed by `increase`, `decrease`, or `affect`.
- `geneName`: Gene name if provided, otherwise leave blank.
- `geneSymbol`: Gene symbol if provided, otherwise leave blank.

#### Guidelines:
1. Extract specific relationships between these chemicals and genes using only the predefined terms from the provided relationship list. Do not use any terms or descriptions of interactions from the source text unless they exactly match terms from the list.
2. Provide only the CSV output—no additional explanations or comments.
3. Ensure that your CSV output is well-formed and follows CSV format.
4. Recheck to ensure the relationship is in the provided relationship list.
5. **Determining increase vs. decrease in expression—use ALL available context:**
   - **If sheet name is provided:** Look for keywords like "up", "down", "increase", "decrease", "upregulated", "downregulated" in the sheet name.
   - **If sheet name is not provided or unclear:** Use the table caption and column headers:
     - Look for phrases like "up-regulated in [chemical] treatment" → chemical INCREASES expression
     - Look for phrases like "down-regulated in [chemical] treatment" → chemical DECREASES expression
     - Look for comparison direction in column headers
   - **Fold change interpretation (when present):**
     - Positive fold change typically means first condition > second condition
     - Negative fold change (or negative log2) typically means first condition < second condition
     - Always check column headers to understand what is being compared (e.g., "Control vs Treatment" or "Treatment vs Control")
   - **If directionality is unclear:** Use "affect expression" instead of "increase expression" or "decrease expression"

—
### Example 1: With Sheet Name

#### Table Caption:
Genes with significant changes in mRNA expression levels after bisphenol A treatment

#### Table Sheet Name:
The control decreased by more than 1.

#### Example Table:

```
ID,Gene Symbol,Description,log2 fold change
1,BRCA1,BReast CAncer gene 1,-2.7
2,TP53,tumor protein p53,-2.1
```

#### Example CSV Output:

```
chemicalName,relationship,geneName,geneSymbol
"bisphenol A","increase expression","BReast CAncer gene 1","BRCA1"
"bisphenol A","increase expression","tumor protein p53","TP53"
```

#### Logic: Sheet name indicates genes DOWN in control, so bisphenol A INCREASES expression.

—
### Example 2: Without Sheet Name (using caption)

#### Table Caption:
Genes up-regulated at least 1.5-fold in bisphenol A treatment in rats

#### Table Sheet Name:
Not provided

#### Example Table:

```
ID,Description,intensity, treatment
1, BReast CAncer gene 1,350,5000
2,tumor protein p53,200,500
```

#### Example CSV Output:

```
chemicalName,relationship,geneName,geneSymbol
"bisphenol A","increase expression","BReast CAncer gene 1",""
"bisphenol A","increase expression","tumor protein p53",""
```

#### Logic: Caption says "up-regulated in bisphenol A treatment" → bisphenol A INCREASES expression.

—
### Example 3: Ambiguous Direction

#### Table Caption:
Genes with altered expression after treatment

#### Table Sheet Name:
Not provided

#### Example Table:

```
Gene,Change
BRCA1,altered
TP53,modified
```

#### Example CSV Output:

```
chemicalName,relationship,geneName,geneSymbol
"treatment compound","affect expression","BRCA1",""
"treatment compound","affect expression","TP53",""
```

#### Logic: No clear directional information → use "affect expression".
