## Supplementary material for "CLOVER: Chemical-gene LLM benchmark for obtaining valid entity relationships for the Comparative Toxicogenomics Database (CTD)": User prompt for text extraction

### Structured Extraction Prompt for Text using One-Shot Learning

#### Task: Extract Chemical-Gene Relationships

Extract specific relationships between chemicals and genes from the given text. Format the output as CSV with one relationship per row.

##### Input Text:
The alkaline phosphatase activity reached a maximum value on day 3 of the secondary culture. Simvastatin increased the alkaline phosphatase activity in a dose dependent manner, and the increase in the activity was significant at a statin concentration of $10^{-6}$ M at 3 ($p<0.01$) and 6 days ($p<0.05$) in the culture compared to the control cultures. The stimulating effect of simvastatin was not showed during the late period of the secondary cultures (Fig. 4).

##### Chemicals (Name, MESH ID):
simvastatin, D019821

##### Genes (Name, NCBI ID):
alkaline phosphatase, 250

—

#### Predefined Relationship Types

You must use ONLY the following relationship types. Do not use any other terms:

**Base relationships:**
List of relationships from Fig 5

**Directional prefixes:**
These relationships can be prefixed with one of the following to indicate direction:

- `increase`—when the text indicates upregulation, enhancement, stimulation, or increase
- `decrease`—when the text indicates downregulation, inhibition, reduction, or decrease
- `affect`—when the text indicates a change or effect but the direction is unclear or mixed

**Examples of valid relationships:**

- "increase expression"
- "decrease activity"
- "affect phosphorylation"
- "binding" (no prefix needed for non-directional relationships)

—

#### Output Format

Generate a CSV with the following columns (include the header row):

**explanation, chemicalName, chemicalID, relationship, geneName, geneID, source**

**Field Descriptions:**

1. **explanation**: Brief description of the interaction (1–2 sentences max)
2. **chemicalName**: Exact name of the chemical from the provided list
3. **chemicalID**: MESH ID of the chemical
4. **relationship**: Relationship type from the predefined list (with prefix if applicable)
5. **geneName**: Exact name of the gene from the provided list
6. **geneID**: NCBI ID of the gene
7. **source**: The exact verbatim text snippet from the input that supports this relationship (use quotation marks if the snippet contains commas)

—

#### Extraction Guidelines

**DO:**

1. Extract only relationships explicitly stated or clearly implied in the text
2. Use only the predefined relationship types from the list above
3. Map common verbs to the appropriate relationship:
    - "upregulates", "enhances", "stimulates" → `increase`
    - "downregulates", "inhibits", "suppresses", "reduces" → `decrease`
    - "modulates", "alters", "changes", "affects" → `affect`
4. Extract each distinct chemical-gene pair as a separate row
5. Use the exact chemical and gene names from the provided lists
6. Ensure all CSV fields are properly formatted and escaped

**DO NOT:**

1. Make assumptions about relationships not stated in the text
2. Use relationship terms not in the predefined list
3. Add interpretations or additional context beyond what is in the text
4. Extract vague or uncertain relationships
5. Include explanatory text outside the CSV format
6. Invent or paraphrase the source text—it must be verbatim
7. Extract negative relationships (e.g., "chemical X does NOT affect gene Y", "no effect on", "did not change")
8. Extract null findings or lack of effect—only extract positive findings where an interaction exists

—

#### Example

##### Example Input Text:
"Exposure to cadmium chloride increases expression of MMP13 mRNA and MMP9 mRNA in cartilage cells. Bisphenol A reduces the activity of TOP2A protein and upregulates TP53 expression in chondrocytes. Treatment with lead acetate did not affect COL2A1 expression."

##### Example Output:

```
explanation,chemicalName,chemicalID,relationship,geneName,geneID,source
"cadmium chloride increases expression of MMP13","cadmium chloride","D019256",
"increase expression","MMP13","4332","Exposure to cadmium chloride increases expression
of MMP13 mRNA and MMP9 mRNA in cartilage cells."
"cadmium chloride increases expression of MMP9","cadmium chloride","D019256",
"increase expression","MMP9","4318","Exposure to cadmium chloride increases expression
of MMP13 mRNA and MMP9 mRNA in cartilage cells."
"bisphenol A decreases activity of TOP2A","bisphenol A","D001778",
"decrease activity","TOP2A","7153","Bisphenol A reduces the activity of TOP2A protein
and upregulates TP53 expression in chondrocytes."
"bisphenol A increases expression of TP53","bisphenol A","D001778",
"increase expression","TP53","7157","Bisphenol A reduces the activity of TOP2A
protein and upregulates TP53 expression in chondrocytes."
```

**Note:** The relationship "lead acetate did not affect COL2A1 expression" is NOT extracted because it describes a null finding (no effect).

—

#### Your Task
Now extract all chemical-gene relationships from the input text provided above following these guidelines. Output only the CSV format with no additional commentary.
