## Supplementary material for "CLOVER: Chemical-gene LLM benchmark for obtaining valid entity relationships for the Comparative Toxicogenomics Database (CTD)": User prompt for figure validation

### Structured Validation Prompt for Figures using One-Shot Learning

Validate and correct relationships between chemicals and genes extracted from a figure/image.

—

#### Validation Rules

1. **Relationship Type:** Must match the predefined list. Replace invalid relationships with the closest valid alternative supported by the image; mark for removal (`agreement=0`) if no valid alternative exists. Chemical-only relationships (`abundance`, `response to chemical`, `chemical synthesis`) are invalid for chemical–gene pairs.
2. **Prefix Rules:** All relationships require a directional prefix (`increase`, `decrease`, `affect`) except `cotreatment` and `binding`. Ensure consistent lowercase capitalization.
3. **Image Direction Indicators:** Use visual cues to determine directionality — taller/darker bars, upward slopes, ↑ symbols, and warm colors indicate increase; shorter/lighter bars, downward slopes, ↓ symbols, and cool colors indicate decrease.
4. **Entity Validation:** Each relationship must involve exactly two entities. Use "-" for missing gene name or symbol. Set `agreement=0` if entity information is incomplete.
5. **Null Relationships:** Remove interactions marked "NS" or "n.s.", showing no visible difference, or indicating no effect.

—

#### Predefined Relationships:

List of relationships from Fig 5

—

#### Input:

- **Image Caption:** Cytotoxicity of SM on SAE and BTE cells. Cells were exposed to various concentrations of SM. Cell viability was measured using the MTS assay 24 h after exposure, and is expressed as a percentage of the control (unexposed cells, "C"). *** P < 0.001 versus control.
- **Extracted Relationships:**

```
chemicalName,relationship,geneName,geneSymbol
"Sulfur Mustard","Decreases abundance","-","-"
```

—

#### Output Format

Return CSV with header: `chemicalName, relationship, geneName, geneSymbol, agreement, reasoning`

- `agreement`: 1 (correct as-is) or 0 (needs correction/removal)
- `reasoning`: blank if `agreement=1`, otherwise explain what changed and why

—

#### Example

**Input:**

```
chemicalName,relationship,geneName,geneSymbol
"Genistein","Decreases expression","-","DEF2"
"Genistein","Increases expression","-","ABC1"
"Genistein","activity","tyrosine 3-monooxygenase","-"
```

**Caption:** Gene regulation by Genistein.

**Output:**

```
chemicalName,relationship,geneName,geneSymbol,agreement,reasoning
"Genistein","decrease expression","-","DEF2","0",
"Fixed capitalization: 'Decreases' to 'decrease'"
"Genistein","increase expression","-","ABC1","0",
"Fixed capitalization: 'Increases' to 'increase'"
"Genistein","affect activity","tyrosine 3-monooxygenase","-","0",
"Added 'affect' prefix: 'activity' requires a directional prefix"
```

—

Validate each row. Output only the CSV, no additional commentary.
