## Supplementary material for "CLOVER: Chemical-gene LLM benchmark for obtaining valid entity relationships for the Comparative Toxicogenomics Database (CTD)": User prompt for table validation.

**Structured Validation Prompt for Tables using One-Shot Learning**

Validate and correct relationships between chemicals and genes extracted from a table.

—

### Validation Rules

1. **Relationship Type:** Must match the predefined list. Replace invalid relationships with the closest valid alternative supported by the table; mark for removal (`agreement=0`) if no valid alternative exists. Chemical-only relationships (`abundance`, `response to chemical`, `chemical synthesis`) are invalid for chemical–gene pairs.
2. **Prefix Rules:** All relationships require a directional prefix (`increase`, `decrease`, `affect`) except `cotreatment` and `binding`. Use table context to determine direction: positive values, ↑, or fold-change >1.0 indicate increase; negative values, ↓, or fold-change <1.0 indicate decrease.
3. **Entity Validation:** Each relationship must involve exactly two entities. Set `agreement=0` if entity information is missing or incomplete.
4. **Null Relationships:** Remove interactions marked "NS", "no change", "0.00", empty effect cells, or fold-change = 1.0 with NS notation.
5. **Table Context:** Verify that the relationship type and prefix direction align with column headers, table symbols, and fold-change values.

—

### Predefined Relationships:

List of relationships from Fig 5

—

### Input:

- **Table Caption:** High level functions most significantly associated with indomethacin-induced gene expression profile.
- **Table XML:** XML table content
- **Extracted Relationships:**

```
chemicalName,chemicalID,relationship,geneName,geneID
"indomethacin","D007213","increase expression","G6PD","2539"
"indomethacin","D007213","increase expression","GPX1","2876"
...
"indomethacin","D007213","increase expression","MADA","-"
```

—

### Output Format

Return CSV with header: `chemicalName, chemicalID, relationship, geneName, geneID, agreement, reasoning`

- `agreement`: 1 (correct as-is) or 0 (needs correction/removal)
- `reasoning`: blank if `agreement=1`, otherwise explain what changed and why

—

### Example

**Input:**

```
chemicalName,chemicalID,relationship,geneName,geneID
"cadmium chloride","D019256","increase levels","MMP13","4332"
"Lead acetate","D007854","affect expression","COL2A1","1280"
```

**Table shows:** Cadmium increases expression 2.3-fold; Lead shows 1.0 (NS).

**Output:**

```
chemicalName,chemicalID,relationship,geneName,geneID,agreement,reasoning
"cadmium chloride","D019256","increase expression","MMP13","4332","0",
"Changed 'increase levels' to 'increase expression':
'levels' not in predefined list"
"Lead acetate","D007854","affect expression","COL2A1","1280","0",
"Null relationship: table shows 1.0 (NS), no significant change"
```

—

Validate each row. Output only the CSV, no additional commentary.
