## Supplementary material for "CLOVER: Chemical-gene LLM benchmark for obtaining valid entity relationships for the Comparative Toxicogenomics Database (CTD)": User prompt for supplement validation

### Structured Validation Prompt for CSV Supplements using One-Shot Learning

Validate and correct relationships between chemicals and genes extracted from a supplemental table.

—

#### Validation Rules

1. **Relationship Type:** Must match the predefined list. Replace invalid relationships with the closest valid alternative supported by the table; mark for removal (`agreement=0`) if no valid alternative exists. Chemical-only relationships (`abundance`, `response to chemical`, `chemical synthesis`) are invalid for chemical–gene pairs.
2. **Prefix Rules:** All relationships require a directional prefix (`increase`, `decrease`, `affect`) except `cotreatment` and `binding`. Use `affect` when direction is unclear.
3. **Direction Validation:** Verify prefix direction using all available context:
   - **Sheet name:** "down" or "decrease" in sheet name → genes DOWN in control → chemical INCREASES expression; "up" or "increase" → chemical DECREASES expression.
   - **Caption/headers:** "up-regulated in [chemical]" → increase; "down-regulated in [chemical]" → decrease.
   - **Fold-change:** >1.0 or positive log2 = increase; <1.0 or negative log2 = decrease (verify comparison direction from column headers).
4. **Entity Validation:** Each relationship must involve exactly two entities. Set `agreement=0` if entity information is missing or incomplete.
5. **Null Relationships:** Remove interactions marked "NS", "no change", fold-change = 1.0, or ∼0 in log scale.

—

#### Predefined Relationships:

List of relationships from Fig 5

—

#### Input:

- **Table Caption:** Genes with a time-dependent, continuous increase or decrease in mRNA expression level during bisphenol A treatment.
- **Sheet Name:** ctr up >1.5
- **Table CSV:** CSV supplement content
- **Extracted Relationships:**

```
chemicalName,relationship,geneName,geneSymbol
"bisphenol A","decrease expression",
    "acetyl-Coenzyme A acetyltransferase 2","ACAT2"
...
"bisphenol A","decrease expression",
    "low density lipoprotein receptor-related protein 8","LRP8"
```

—

#### Output Format

Return CSV with header: `chemicalName, relationship, geneName, geneSymbol, agreement, reasoning`

- `agreement`: 1 (correct as-is) or 0 (needs correction/removal)
- `reasoning`: blank if `agreement=1`, otherwise explain what changed and why

—

#### Example

**Caption:** Genes with significant changes after bisphenol A treatment.
**Sheet Name:** Control decreased.

**Input:**

```
chemicalName,relationship,geneName,geneSymbol
"bisphenol A","decrease expression","carbonyl reductase 1","CR1"
"bisphenol A","increase expression",
    "glucose-6-phosphate dehydrogenase","G6PD"
```

**Output:**

```
chemicalName,relationship,geneName,geneSymbol,agreement,reasoning
"bisphenol A","increase expression","carbonyl reductase 1","CR1","0",
"Changed 'decrease' to 'increase' sheet name indicates
genes down in control",
"bisphenol A","increase expression",
"glucose-6-phosphate dehydrogenase","G6PD","1",""
```

—

Validate each row. Output only the CSV, no additional commentary.
