## Supplementary material for "CLOVER: Chemical-gene LLM benchmark for obtaining valid entity relationships for the Comparative Toxicogenomics Database (CTD)": User prompt for text validation.

### Structured Validation Prompt for Text using One-Shot Learning

Validate and correct relationships between chemicals and genes using ONLY the information provided below.

—

#### Validation Rules

1. **Relationship Type:** Must match the predefined list. Replace invalid relationships with the closest valid alternative supported by the text; mark for removal (`agreement=0`) if no valid alternative exists. Chemical-only relationships (`abundance`, `response to chemical`, `chemical synthesis`) are invalid for chemical–gene pairs.
2. **Prefix Rules:** All relationships require a directional prefix (`increase`, `decrease`, `affect`) except `cotreatment` and `binding`. Use `affect` only when direction cannot be determined from the text. Remove any prefix applied to exempt relationships.
3. **Entity Validation:** Each relationship must involve exactly two entities. Set `agreement=0` if entity information is missing or incomplete; do not guess missing entities.
4. **Null Relationships:** Remove interactions where the source text indicates no effect (e.g., "did not change", "no effect on", "failed to alter").
5. **Prior Work:** Remove interactions cited from previous studies or other works (e.g., "previous studies showed", "it has been reported", "as demonstrated by").

—

#### Predefined Relationships:

List of relationships from Fig 5

—

#### Input:

- **Original Text:** Section passage
- **Extracted Relationships:**

```
explanation,chemicalName,chemicalID,relationship,geneName,
geneID,source
"MTT assay used to measure cellular growth in cells treated for
alkaline phosphatase activity assay","MTT","C070243","activity",
"alkaline phosphatase","250","The cells were seeded in 6 well
plates and treated in the same manner as with the alkaline
phosphatase activity assay."
```

—

#### Output Format

Return CSV with header: `explanation, chemicalName, chemicalID, relationship, geneName, geneID, source, agreement, reasoning`

- `agreement`: 1 (correct as-is) or 0 (needs correction/removal)
- `reasoning`: blank if `agreement=1`, otherwise specify which rule was violated and what changed

—

#### Example

**Original Text:** "Exposure to cadmium chloride increases levels of MMP13 and MMP9. Bisphenol A reduces activity of TOP2A and affects expression of TP53 in chondrocytes. Lead acetate did not change COL2A1 expression. Previous studies showed that arsenic increases oxidation of KEAP1."

**Input:**

```
explanation,chemicalName,chemicalID,relationship,geneName,geneID,source
"cadmium chloride increases levels of MMP13","cadmium chloride",
"D019256","increase levels","MMP13","4332",
"Exposure to cadmium chloride increases levels of MMP13 and MMP9."
"cadmium chloride increases expression of MMP9","cadmium chloride",
"D019256","response to chemical","MMP9","4318",
"Exposure to cadmium chloride increases levels of MMP13 and MMP9."
"bisphenol A decreases activity of TOP2A","bisphenol A",
"D001778","decrease activity","TOP2A","7153",
"Bisphenol A reduces activity of TOP2A and affects expression of TP53."
"bisphenol A affects expression of TP53","bisphenol A",
"D001778","affect expression","TP53","7157",
"Bisphenol A reduces activity of TOP2A and affects expression of TP53."
"lead acetate has no effect on COL2A1","lead acetate",
"D007854","affect expression","COL2A1","1280",
"Lead acetate did not change COL2A1 expression."
"arsenic increases oxidation of KEAP1","arsenic",
"D001151","increase oxidation","KEAP1","9817",
"Previous studies showed that arsenic increases oxidation of KEAP1."
```

**Output:**

```
explanation,chemicalName,chemicalID,relationship,geneName,
geneID,source,agreement,reasoning
"cadmium chloride increases expression of MMP13","cadmium chloride",
"D019256","increase expression","MMP13","4332",
"Exposure to cadmium chloride increases levels of MMP13 and MMP9.","0",
"Changed 'increase levels' to 'increase expression':
'levels' is not in the predefined list"
"cadmium chloride increases expression of MMP9","cadmium chloride",
"D019256","increase expression","MMP9","4318",
"Exposure to cadmium chloride increases levels of MMP13 and MMP9.","0",
"Changed 'response to chemical' to 'increase expression':
chemical-only relationship invalid for chemical-gene pair"
"bisphenol A decreases activity of TOP2A","bisphenol A",
"D001778","decrease activity","TOP2A","7153",
"Bisphenol A reduces activity of TOP2A and affects expression of TP53.","1",""
"bisphenol A affects expression of TP53","bisphenol A",
"D001778","affect expression","TP53","7157",
"Bisphenol A reduces activity of TOP2A and affects expression of TP53.","1",""
"lead acetate has no effect on COL2A1","lead acetate",
"D007854","affect expression","COL2A1","1280",
"Lead acetate did not change COL2A1 expression.","0",
"Null relationship: source text states 'did not change'"
"arsenic increases oxidation of KEAP1","arsenic",
"D001151","increase oxidation","KEAP1","9817",
"Previous studies showed that arsenic increases oxidation of KEAP1.","0",
"Relationship from previous studies/other works"
```

—

Validate each row. Output only the CSV, no additional commentary.
